# Cohesin mutations are synthetic lethal with stimulation of WNT signaling

**DOI:** 10.1101/2020.07.23.218875

**Authors:** Chue Vin Chin, Jisha Antony, Sarada Ketharnathan, Gregory Gimenez, Kate M. Parsons, Jinshu He, Amee J. George, Antony Braithwaite, Parry Guilford, Ross D. Hannan, Julia A. Horsfield

## Abstract

Mutations in genes encoding subunits of the cohesin complex are common in several cancers, but may also expose druggable vulnerabilities. We generated isogenic MCF10A cell lines with deletion mutations of genes encoding cohesin subunits SMC3, RAD21 and STAG2 and screened for synthetic lethality with 3,009 FDA-approved compounds. The screen identified several compounds that interfere with transcription, DNA damage repair and the cell cycle. Unexpectedly, one of the top ‘hits’ was a GSK3 inhibitor, an agonist of Wnt signaling. We show that sensitivity to GSK3 inhibition is likely due to stabilization of β-catenin in cohesin mutant cells, and that Wnt-responsive gene expression is highly sensitized in *STAG2*-mutant CMK leukemia cells. Moreover, Wnt activity is enhanced in zebrafish mutant for cohesin subunit *rad21*. Our results suggest that cohesin mutations could progress oncogenesis by enhancing Wnt signaling, and that targeting the Wnt pathway may represent a novel therapeutic strategy for cohesin mutant cancers.

## Introduction

The cohesin complex is essential for sister chromatid cohesion, DNA replication, DNA repair, and genome organization. Three subunits, SMC1A, SMC3 and RAD21, form the core ring-shaped structure of human cohesin [1, 2]. A fourth subunit of either STAG1 or STAG2 binds to cohesin by contacting RAD21 and SMC subunits [3], and is required for the association of cohesin with DNA [1-3]. The STAG subunits of cohesin are also capable of binding RNA in the nucleus [4]. Cohesin associates with DNA by interaction with loading factors NIPBL and MAU2 [5], its stability on DNA is regulated by the acetyltransferases ESCO1 [6] and ESCO2 [7], and its removal is facilitated by PDS5 and WAPL [8, 9]. The cohesin ring itself acts as a molecular motor to extrude DNA loops, and this activity is thought to underlie its ability to organize the genome [10, 11]. Cohesin works together with CCCTC-binding factor (CTCF) to mediate three-dimensional genome structures, including enhancer-promoter loops that instruct gene accessibility and expression [12, 13]. The consequences of cohesin mutation therefore manifest as chromosome segregation errors, DNA damage, and deficiencies in genome organization leading to gene expression changes.

All cohesin subunits are essential to life: homozygous mutations in genes encoding complex members are embryonic lethal [14]. However, haploinsufficient germline mutations in *NIPBL, ESCO2*, and in cohesin genes, cause human developmental syndromes known as the “cohesinopathies” [14]. Somatic mutations in cohesin genes are prevalent in several different types of cancer, including bladder cancer (15-40%), endometrial cancer (19%), glioblastoma (7%), Ewing’s sarcoma (16-22%) and myeloid leukemias (5-53%) [15-17]. The prevalence of cohesin gene mutations in myeloid malignancies [18-22] reflects cohesin’s role in determining lineage identity and differentiation of hematopoietic cells [23-26]. Of the cohesin genes, *STAG2* is the most frequently mutated, with about half of cohesin mutations in cancer involving *STAG2* [17].

While cancer-associated mutations in genes encoding RAD21, SMC3 and STAG1 are always heterozygous [21, 27, 28], mutations in the X chromosome-located genes *SMC1A* and *STAG2* can result in complete loss of function due to hemizygosity (males), or silencing of the wild type during X-inactivation (females). STAG2 and STAG1 have redundant roles in cell division, therefore complete loss of STAG2 is tolerated due to partial compensation by STAG1. Loss of both STAG2 and STAG1 leads to lethality [29, 30]. STAG1 inhibition in cancer cells with STAG2 mutation causes chromosome segregation defects and subsequent lethality [31]. Therefore, although partial depletion of cohesin can confer a selective advantage to cancer cells, a complete block of cohesin function will cause cell death. The multiple roles of cohesin provides an opportunity to inhibit the growth of cohesin mutant cancer cells via chemical interference with pathways that depend on normal cohesin function. For example, poly ADP-ribose polymerase (PARP) inhibitors were previously shown to exhibit synthetic lethality with cohesin mutations [17, 31-34]. PARP inhibitors prevent DNA double-strand break repair [35], a process that also relies on cohesin function.

To date, only a limited number of compounds have been identified as inhibitors of cohesin-mutant cells [17]. Here, we sought to identify additional compounds of interest by screening libraries of FDA-approved molecules against isogenic MCF10A cells with deficiencies in RAD21, SMC3 or STAG2. Unexpectedly, our screen identified a novel sensitivity of cohesin-deficient cells to a GSK3 inhibitor that acts as an agonist of the Wnt signaling pathway. We found that β-catenin stabilization upon cohesin deficiency likely contributes to an acute sensitivity of Wnt target genes. The results raise the possibility that sensitization to Wnt signaling in cohesin mutant cells may participate in oncogenesis, and suggest that Wnt agonism could be therapeutically useful for cohesin mutant cancers.

## Results

### Cohesin gene deletion in MCF10A cells results in minor cell cycle defects

To avoid any complications with pre-existing oncogenic mutations, we chose the relatively ‘normal’ MCF10A line for creation and screening of isogenic deletion clones of cohesin genes *SMC3, RAD21*, and *STAG2* (Supplementary Fig. 1a). MCF10A is a near-diploid, immortalized, breast epithelial cell line that exhibits normal epithelial characteristics [36] and has been successfully used for siRNA and small molecule screening [37]. We isolated several *RAD21* and *SMC3* deletion clones, and selected single clones for further characterization that grew normally and were essentially heterozygous. In the selected *RAD21* deletion clone, one of three *RAD21* alleles (on chromosome 8, triploid in MCF10A) was confirmed deleted, with one wild type and one undetermined allele (Supplementary Fig. 1c, e-g). In the selected *SMC3* deletion clone, one of two *SMC3* alleles (on chromosome 10) was deleted (Supplementary Fig. 1d, e-g). In the selected *STAG2* deletion clone (Supplementary Fig. 1b), homozygous loss of *STAG2* was determined by the absence of STAG2 mRNA and protein (Supplementary Fig. 1e-g). For convenience here, we named the three cohesin mutant clones *RAD21+/-, SMC3+/-* and *STAG2-/-*.

**Figure 1.**
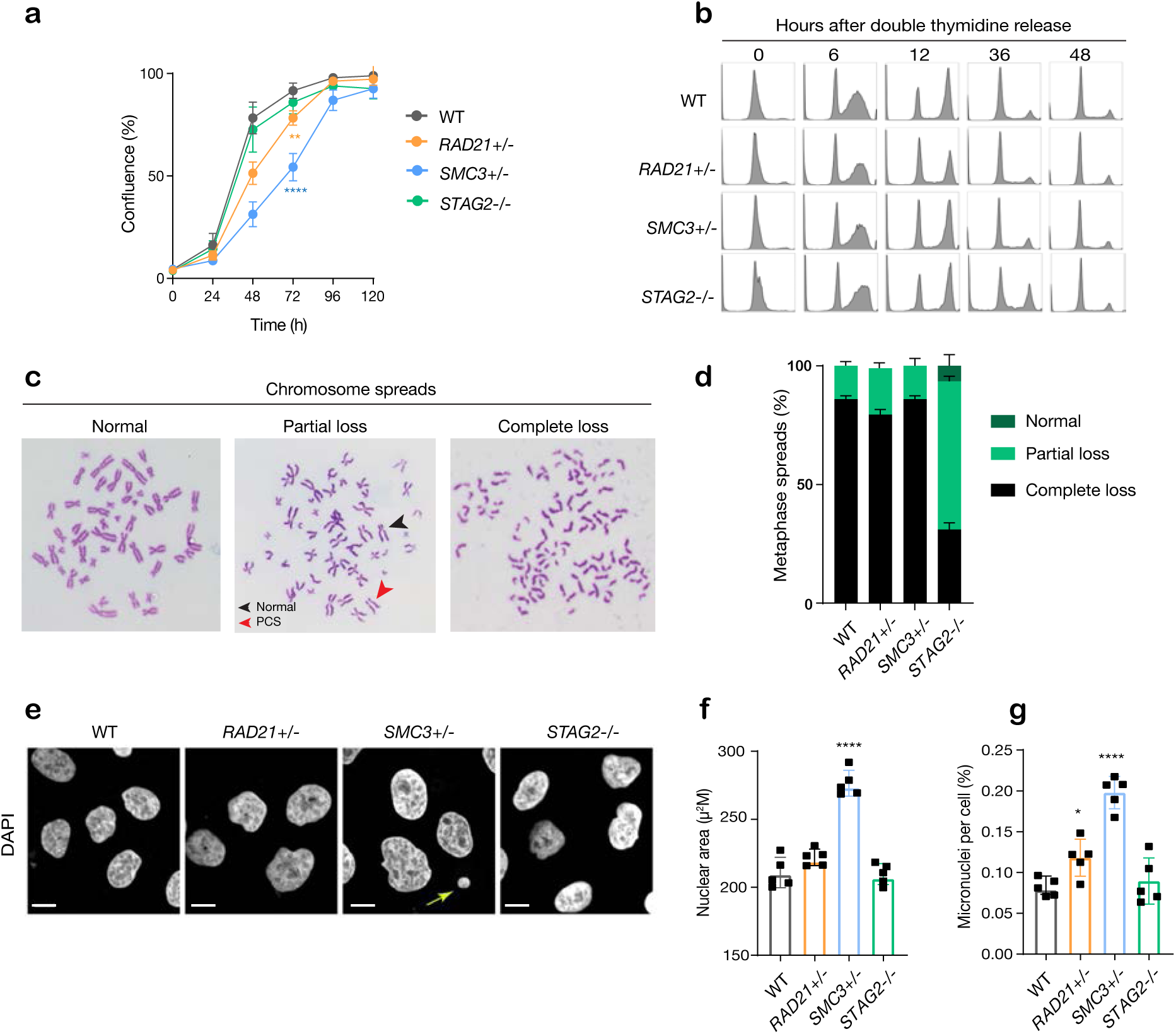
minor deficiencies in cell cycle progression, chromosome segregation and nuclear morphology in cohesin-deficient isogenic cell lines. **a** Proliferation curves (Incucyte) of MCF10A parental and cohesin-deficient clones (n = 3 independent experiments, mean ± sd, two-way ANOVA test: \*\**p* ≤ 0.01; \*\*\*\**p* ≤ 0.0001). Doubling time in hours of MCF10A parental cells are 15.5±0.1; *RAD21+/-*, 17.1±0.2; *SMC3+/-*, 21.4±0.9; *STAG2-/-*, 16.4±0.2 respectively. **b** Flow cytometry analysis of cell cycle progression. **c** Representative metaphase spread images of cohesin-deficient cells. Black arrow indicates a normal chromosome; Red arrow indicates premature chromatid separation (PCS). **d** Quantification of chromosome cohesion defects. A minimum of 20 metaphase spreads were examined per individual experiment. (n=2 independent experiments, mean ± s.d.). **e** Representative confocal images of nuclear morphology. Scale bar, 15 μM. Yellow arrow indicates a micronucleus. **f** Quantification of nuclear area. **g** Quantification of micronuclei (MN). A minimum of 1000 cells was examined per individual experiment. (n = 3 independent experiments, mean ± s.d., \**p* ≤ 0.05; \*\*\*\**p* ≤ 0.0001).

The *RAD21+/-, SMC3+/-* and *STAG2-/-* clones had essentially normal cell cycle progression when compared to parental cells, although the *RAD21*+/-and *SMC3*+/-clones proliferated more slowly than the others (Fig. 1a, b). Chromosome spreads revealed that only the *STAG2*-/-clone had noteworthy chromosome cohesion defects characterized by partial or complete loss of chromosome cohesion and gain or loss of more than one chromosome (Fig. 1c-d, Supplementary Fig. 2a). Remarkably, *SMC3+/-* cells had noticeably larger nuclei that appeared to be less dense (Fig. 1e-f, Supplementary Fig. 2b), possibly owing to decompaction of DNA in this clone. *RAD21+/-* and *SMC3+/-* clones exhibited occasional lagging chromosomes and micronuclei, while the *STAG2-/-* clone did not (Fig. 1g, Supplementary Fig. 2c-e). Cell growth and morphology in monolayer culture was essentially normal in all three cohesin mutant clones (Supplementary Fig. 3).

**Figure 2.**
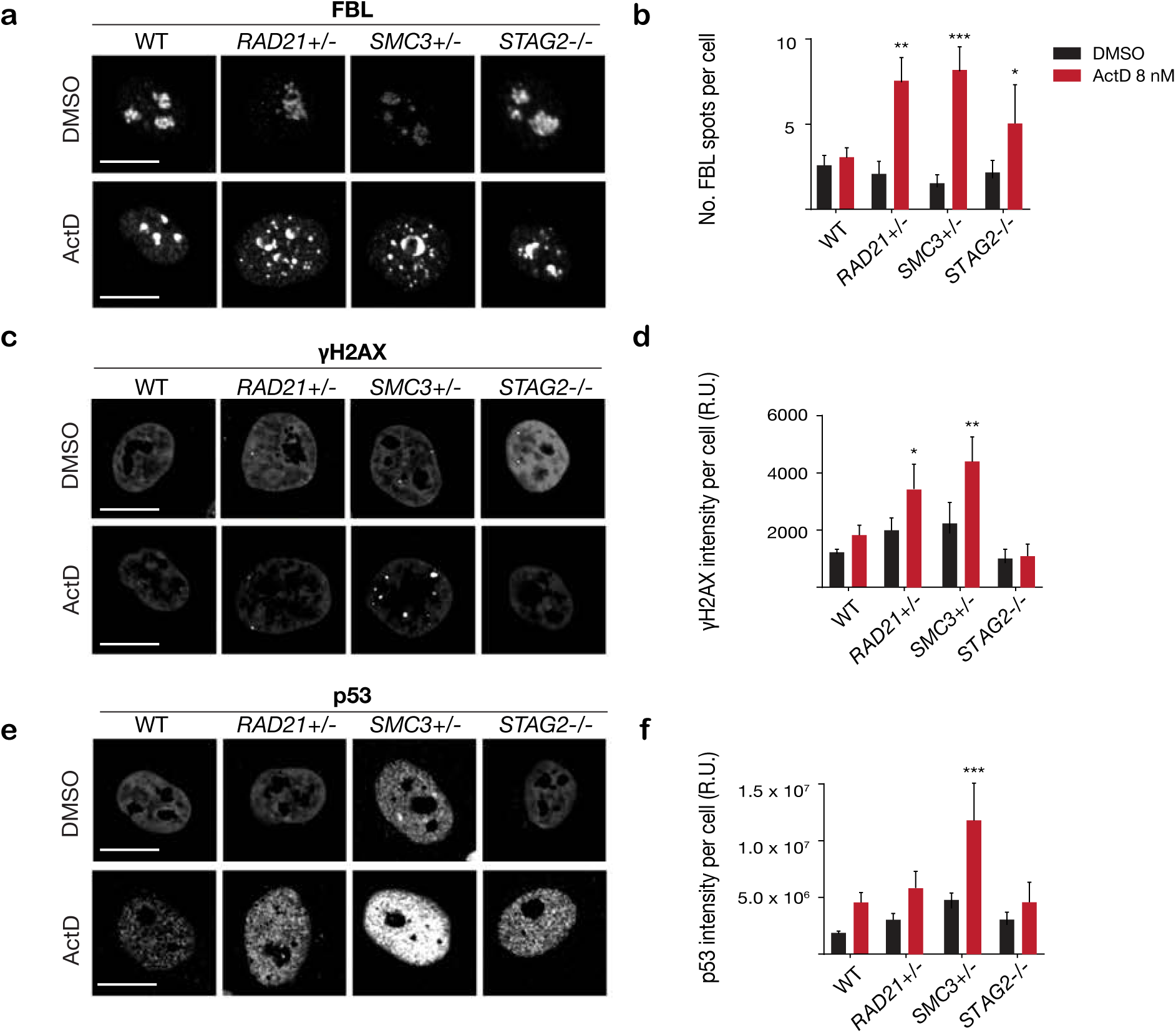
Increased sensitivity to nucleolar stress and DNA damaging agents in cohesin-deficient cells. **a** Representative image and **b** quantification of nucleolar dispersal observed in MCF10A cells after exposure to a DNA damaging agent, Actinomycin D (ActD) 8 nM. Fibrillarin (FBL) staining is used as a marker for nucleoli. **c** Representative image and **d** quantification of DNA damage foci observed in MCF10A cells after exposure to ActD. γH2AX staining is used to detect DNA double-strand breaks; arrows indicate γH2AX foci. **e** Representative image and **f** quantification of nuclear p53 in MCF10A cells after exposure to ActD. A minimum of 500 cells was examined per individual experiment. (n = 3 independent experiments, mean ± s.d., \**p* ≤ 0.05; \*\**p* ≤ 0.01; \*\*\**p* ≤ 0.0005; \*\*\*\**p* ≤ 0.0001). Scale bar, 15 μM

**Figure 3.**
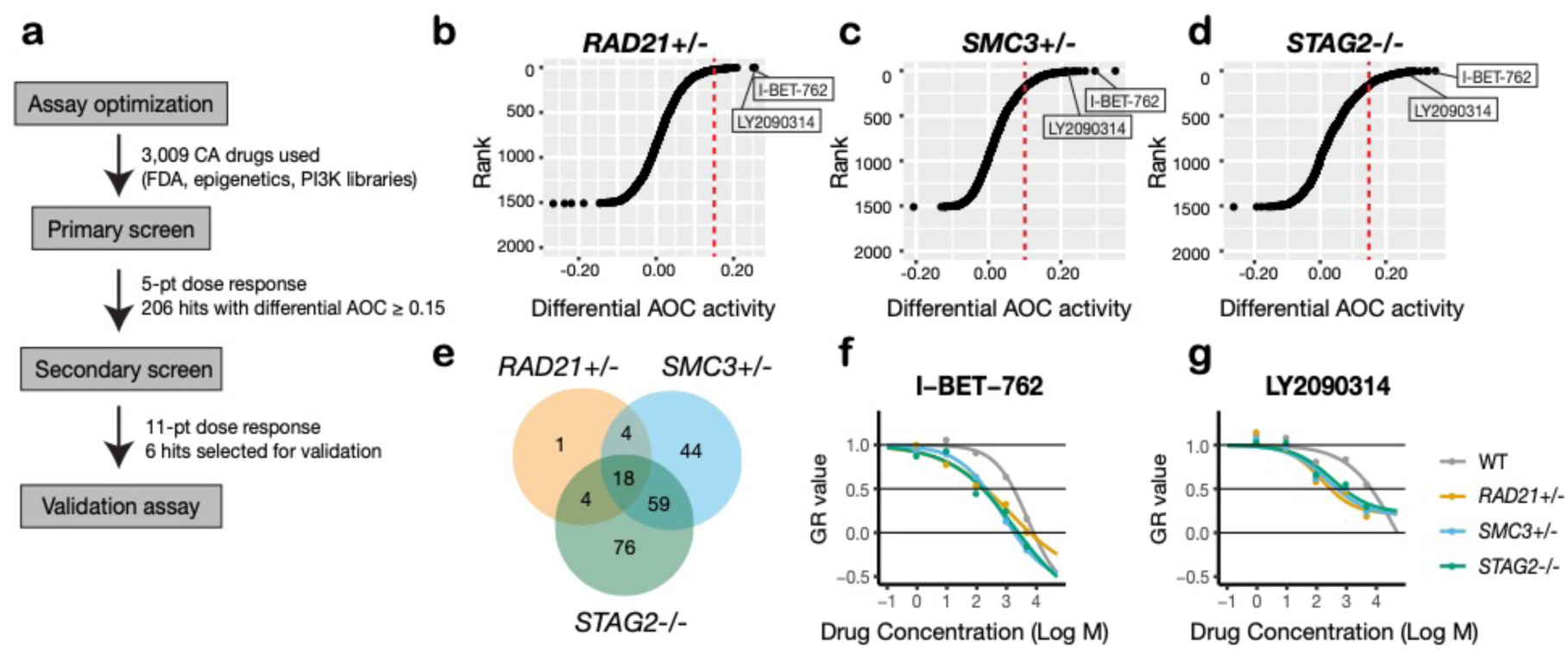
A synthetic lethal screen identifies common sensitivity of cohesin-deficient cells to WNT activation and BET inhibition. **a** Schematic overview of the synthetic lethal screen. **b-d** Overview of the differential area over the curve (AOC) activity of all compounds tested in cohesin-deficient cell lines in the primary screen. A threshold of differential AOC ≥ 0.15 (red dashed lines) was used to filter candidate compounds of interest. **e** Venn diagram showing the number of common and unique compounds that inhibited *RAD21+/-, SMC3+/-* and *STAG2-/-* in the primary screen. **f, g** Dose-response curves of I-BET-762 and LY2090314.

Overall the cohesin deletion clones appear to have infrequent but specific cell cycle anomalies that are shared between some, but not all clones. Anomalies include loss of chromosome cohesion, lagging chromosomes, or micronuclei, but these features do not appear to majorly impact on cell cycle progression or morphology.

### Cohesin-depleted cells have altered nucleolar morphology and are sensitive to DNA damaging agents

Cohesin-deficient cells have been demonstrated to display compromised nucleolar morphology and ribosome biogenesis [38, 39], as well as sensitivity to DNA damaging agents [40, 41]. Analysis of our *RAD21+/-, SMC3+/-* and *STAG2-/-* clones confirmed these findings. Cohesin-deleted cells in steady-state growth had abnormal nucleolar morphology, as revealed by fibrillarin and nucleolin staining (Supplementary Fig. 4). Furthermore, we found that treatment with DNA intercalator/transcription inhibitor Actinomycin D caused marked fragmentation of nucleoli in all three cohesin-deficient cell lines as determined by fibrillarin staining (Fig. 2a, b). Actinomycin D treatment increased γ-H2AX and TP53 in the nuclei of *RAD21+/-* and *SMC3+/-* cells, in particular, indicating that these cells are compromised for DNA damage repair relative to the parental MCF10A cells (Fig. 2c-f). In contrast, immunostaining for γ-H2AX and TP53 levels were comparable at baseline between cohesin-deficient clones and parental cells. Interestingly, the *STAG2-/-* deletion clone was much more resistant to DNA damage caused by Actinomycin D (Fig. 2c-f), even though nucleoli are abnormal in this clone (Supplementary Fig. 4).

**Figure 4.**
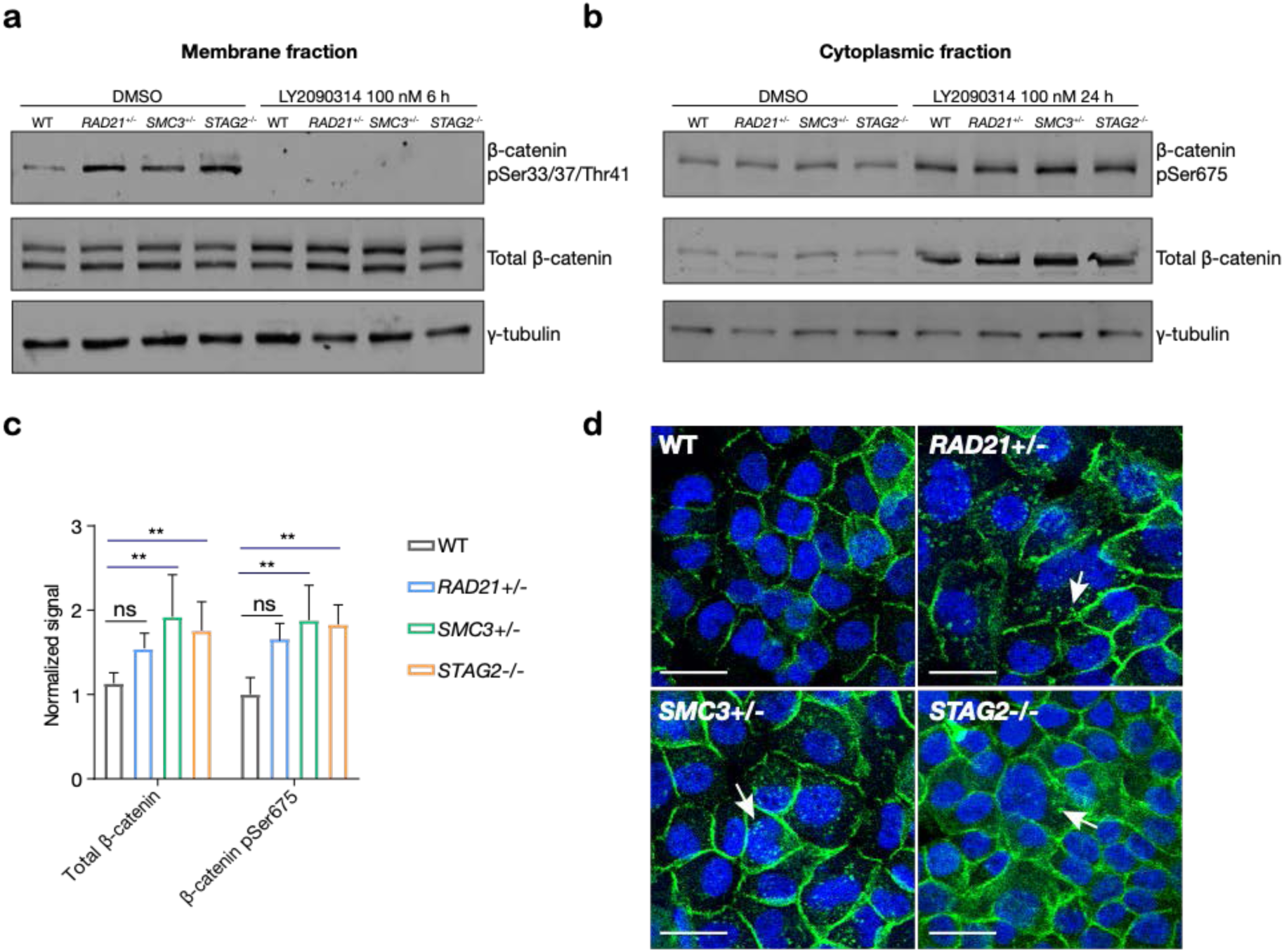
LY2090314-mediated WNT stimulation leads to β-catenin stabilization in cohesin-deficient MCF10A cells. **a** Immunoblot of the membrane fraction of cohesin-deficient MCF10A cells shows increased basal level of β-catenin phosphorylation at Ser33/37/Thr41. **b** Immunoblot of the cytoplasmic fraction shows increased level of both total and phosphorylated β-catenin at Ser675 after MCF10A cells were treated with LY2090314 at 100 nM for 24 hours. **c** Quantification of protein levels for total and phosphorylated beta-catenin at Ser675. (n = 3 independent experiments, mean ± s.d., \*\**p* ≤ 0.01). **d** Immunofluorescence images show cytosolic accumulation of active β-catenin in cohesin-deficient MCF10A cells treated with LY2090314 100 nM for 24 hours. White arrows indicate puncta of β-catenin (Ser675). Scale bar = 25 μM.

Overall, increased susceptibility of *RAD21*+/-and *SMC3*+/-clones to DNA damage is consistent with cohesin’s role in DNA double-strand break repair [42], and highlights the different requirements for RAD21 and SMC3 versus STAG2 in this process. Cohesin mutations were previously shown to sensitize cells to PARP inhibitor, Olaparib [32, 43]. We confirmed a mild to moderate sensitivity to Olaparib in our cohesin-deficient MCF10A clones relative to parental cells (Supplementary Fig. 5a). Collectively, characterization of our cohesin deficient clones provided confidence that they represent suitable models for synthetic lethal screening.

**Figure 5.**
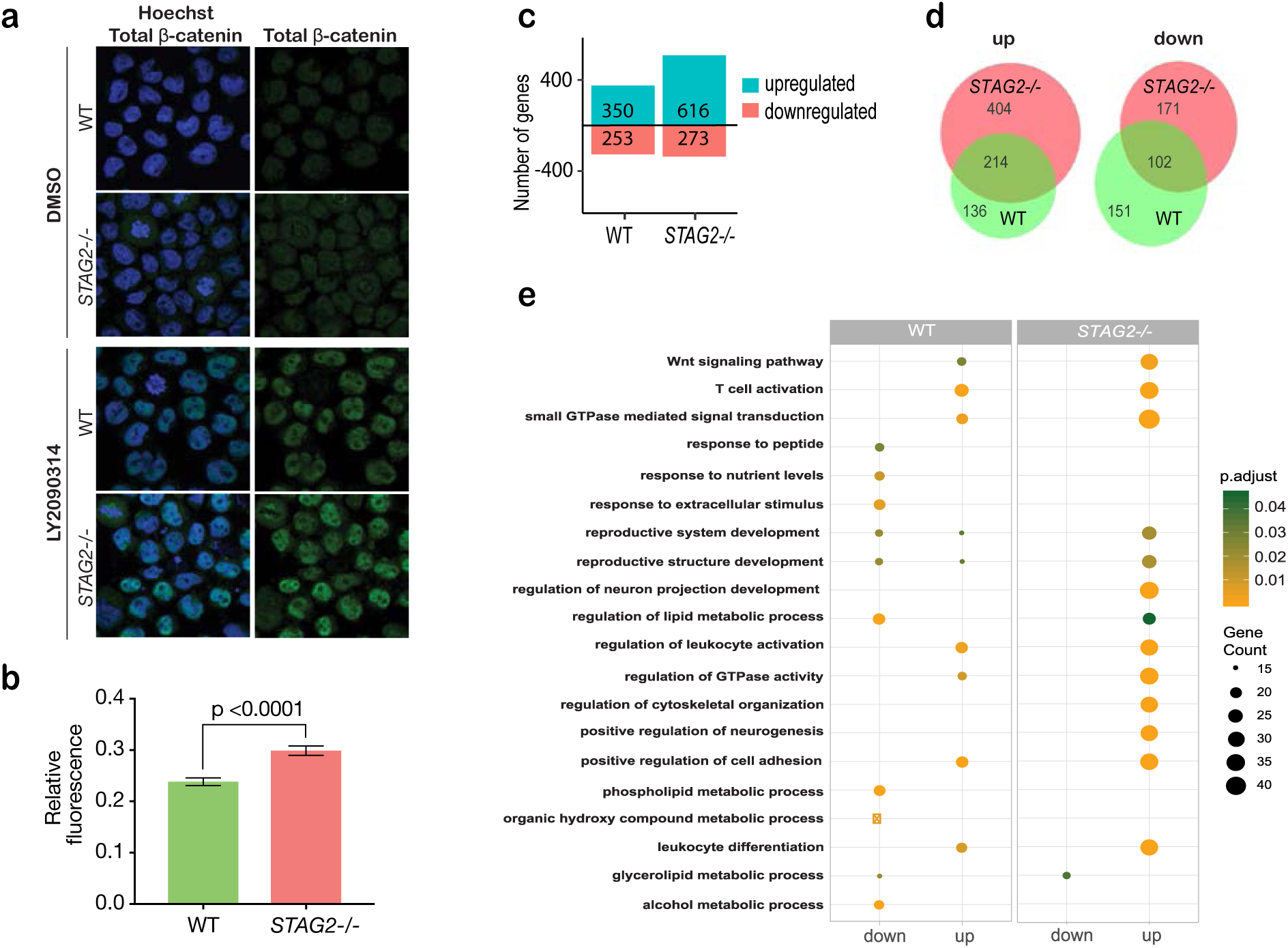
Cohesin-STAG2 mutant CMK cells show increased sensitivity to Wnt signaling. **a** Immunofluorescence images showing slightly increased nuclear accumulation of β-catenin in *STAG2*-CMK cells (STAG2-/-) compared to parental (WT) following treatment with LY2090314 at 100 nM for 24 hours. **b** Quantification of nuclear total β-catenin in parental (WT) and *STAG2*-CMK cells (STAG2-/-) cells. Fluorescence of nuclear total β-catenin was determined relative to the nuclear area. Graph depicts S.E.M. from analyses of 170-188 cell nuclei. **c** Histogram showing the number of genes upregulated or downregulated at FDR ≤ 0.05 upon WNT3A treatment in WT and STAG2-/-CMK cells. **d** Overlap of genes significantly upregulated or downregulated (FDR ≤ 0.05) upon WNT3A treatment between parental and STAG2 mutant cells. **e** Top ten enriched pathways (ranked by gene count) from the significantly downregulated and upregulated genes (FDR ≤ 0.05) following Wnt3a treatment in *STAG2-/-* and parental (WT) CMK cells using the ClusterProfiler R package on Gene Ontology Biological Process dataset modelling for both cell types (WT or *STAG2*-/-) and regulation pattern (up-or downregulation).

### A synthetic lethal screen of FDA-approved compounds identifies common sensitivity of cohesin mutant cells to WNT activation and BET inhibition

To identify additional compounds that inhibit the growth of cohesin-deficient cells, we screened the cohesin-deficient MCF10A cell lines with five dose concentrations (1-5,000 nM) of 3,009 compounds, including FDA-approved compounds (2399/3009), kinase inhibitors (429/3009), and epigenetic modifiers (181/3009) (Fig. 3a). DMSO and Camptothecin were included as negative and positive viability controls, respectively (Supplementary Fig. 5b, c). We assayed cell viability after 48 hours of compound treatment.

Synthetic lethal candidate compounds were ranked based on the differential area over curve (AOC) values that are derived from a growth rate-based metric (GR) (Fig. 3b-d). The GR metric takes into account the varying growth rate of dividing cells to mitigate inconsistent comparison of compound effects across cohesin-deficient cell lines [44]. Compounds that caused ≥30% growth inhibition in cohesin-deficient clones compared with parental MCF10A cells were selected for further analysis. The screen identified candidate 206 synthetic lethal compounds, of which 18 inhibited all three cohesin-deficient cell lines ≥30% more than parental MCF10A cells (Fig. 3e, Table 1, Supplementary Table 1). Most of the 206 compounds inhibited at least two cohesin-deficient cell lines and were classed in similar categories of inhibitor (Supplementary Fig. 6a-d, Supplementary Table 1). Of the 206 primary screen hits, 85 (including common and unique hits identified in each cohesin-deficient clone) were subjected to secondary screening in an 11-point dose curve ranging from 0.5 nM to 10 μM. Notable sensitive pathways included: DNA damage repair, the PI3K/AKT/mTOR pathway, epigenetic control of transcription, and stimulation of the Wnt signaling pathway (Table 1, Supplementary Table 2).

**Table 1.**
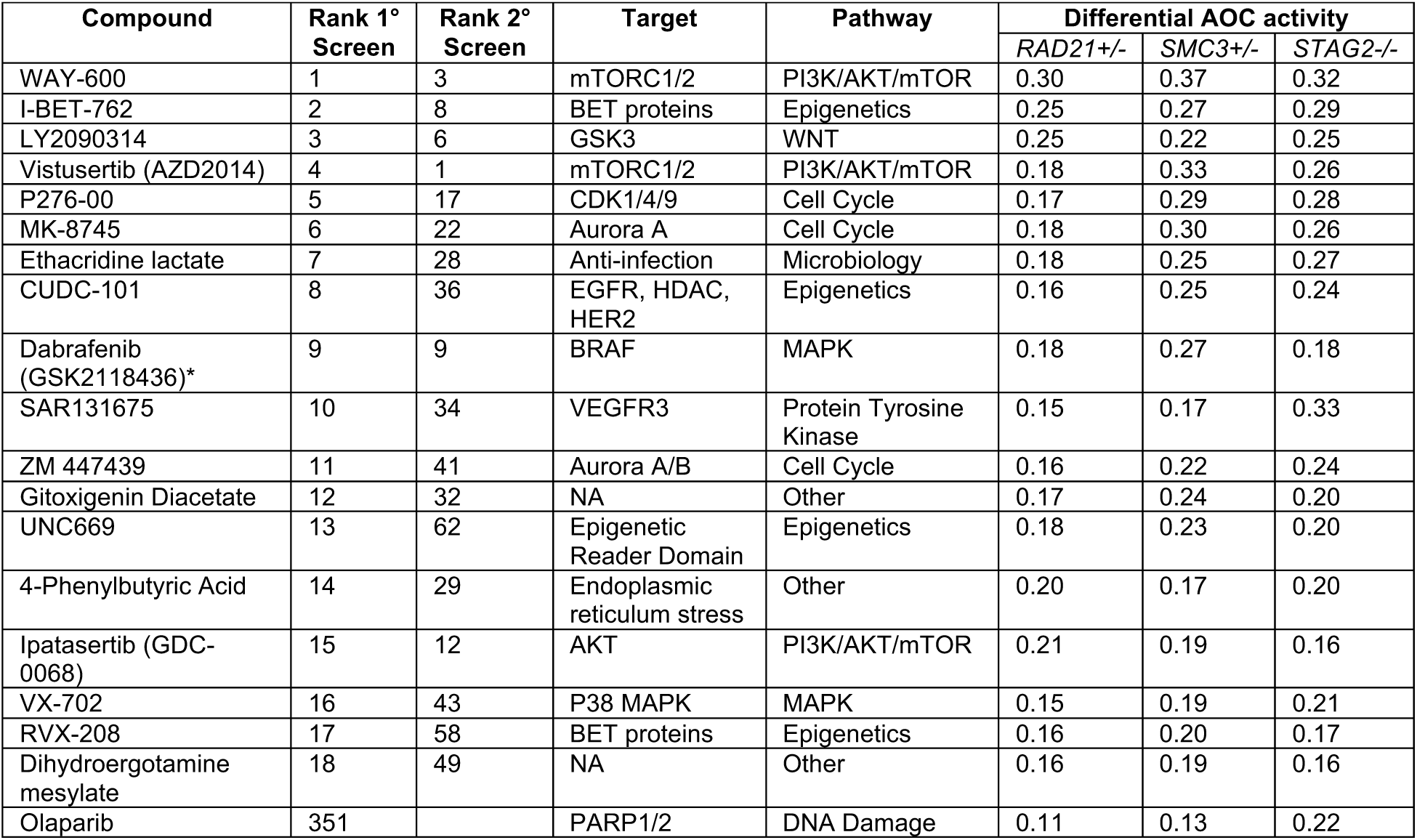
Active compounds identified from the synthetic lethal screen.

| Compound | Rank 1° Screen | Rank 2° Screen | Target | Pathway | Differential AOC activity |  |  |
| --- | --- | --- | --- | --- | --- | --- | --- |
|  |  |  |  |  | *RAD21*+/- | *SMC3*+/- | *STAG2*-/- |
| WAY-600 | 1 | 3 | mTORC1/2 | PI3K/AKT/mTOR | 0.30 | 0.37 | 0.32 |
| I-BET-762 | 2 | 8 | BET proteins | Epigenetics | 0.25 | 0.27 | 0.29 |
| LY2090314 | 3 | 6 | GSK3 | WNT | 0.25 | 0.22 | 0.25 |
| Vistusertib (AZD2014) | 4 | 1 | mTORC1/2 | PI3K/AKT/mTOR | 0.18 | 0.33 | 0.26 |
| P276-00 | 5 | 17 | CDK1/4/9 | Cell Cycle | 0.17 | 0.29 | 0.28 |
| MK-8745 | 6 | 22 | Aurora A | Cell Cycle | 0.18 | 0.30 | 0.26 |
| Ethacridine lactate | 7 | 28 | Anti-infection | Microbiology | 0.18 | 0.25 | 0.27 |
| CUDC-101 | 8 | 36 | EGFR, HDAC, HER2 | Epigenetics | 0.16 | 0.25 | 0.24 |
| Dabrafenib (GSK2118436)* | 9 | 9 | BRAF | MAPK | 0.18 | 0.27 | 0.18 |
| SAR131675 | 10 | 34 | VEGFR3 | Protein Tyrosine Kinase | 0.15 | 0.17 | 0.33 |
| ZM 447439 | 11 | 41 | Aurora A/B | Cell Cycle | 0.16 | 0.22 | 0.24 |
| Gitoxigenin Diacetate | 12 | 32 | NA | Other | 0.17 | 0.24 | 0.20 |
| UNC669 | 13 | 62 | Epigenetic Reader Domain | Epigenetics | 0.18 | 0.23 | 0.20 |
| 4-Phenylbutyric Acid | 14 | 29 | Endoplasmic reticulum stress | Other | 0.20 | 0.17 | 0.20 |
| Ipatasertib (GDC-0068) | 15 | 12 | AKT | PI3K/AKT/mTOR | 0.21 | 0.19 | 0.16 |
| VX-702 | 16 | 43 | P38 MAPK | MAPK | 0.15 | 0.19 | 0.21 |
| RVX-208 | 17 | 58 | BET proteins | Epigenetics | 0.16 | 0.20 | 0.17 |
| Dihydroergotamine mesylate | 18 | 49 | NA | Other | 0.16 | 0.19 | 0.16 |
| Olaparib | 351 |  | PARP1/2 | DNA Damage | 0.11 | 0.13 | 0.22 |

**Figure 6.**
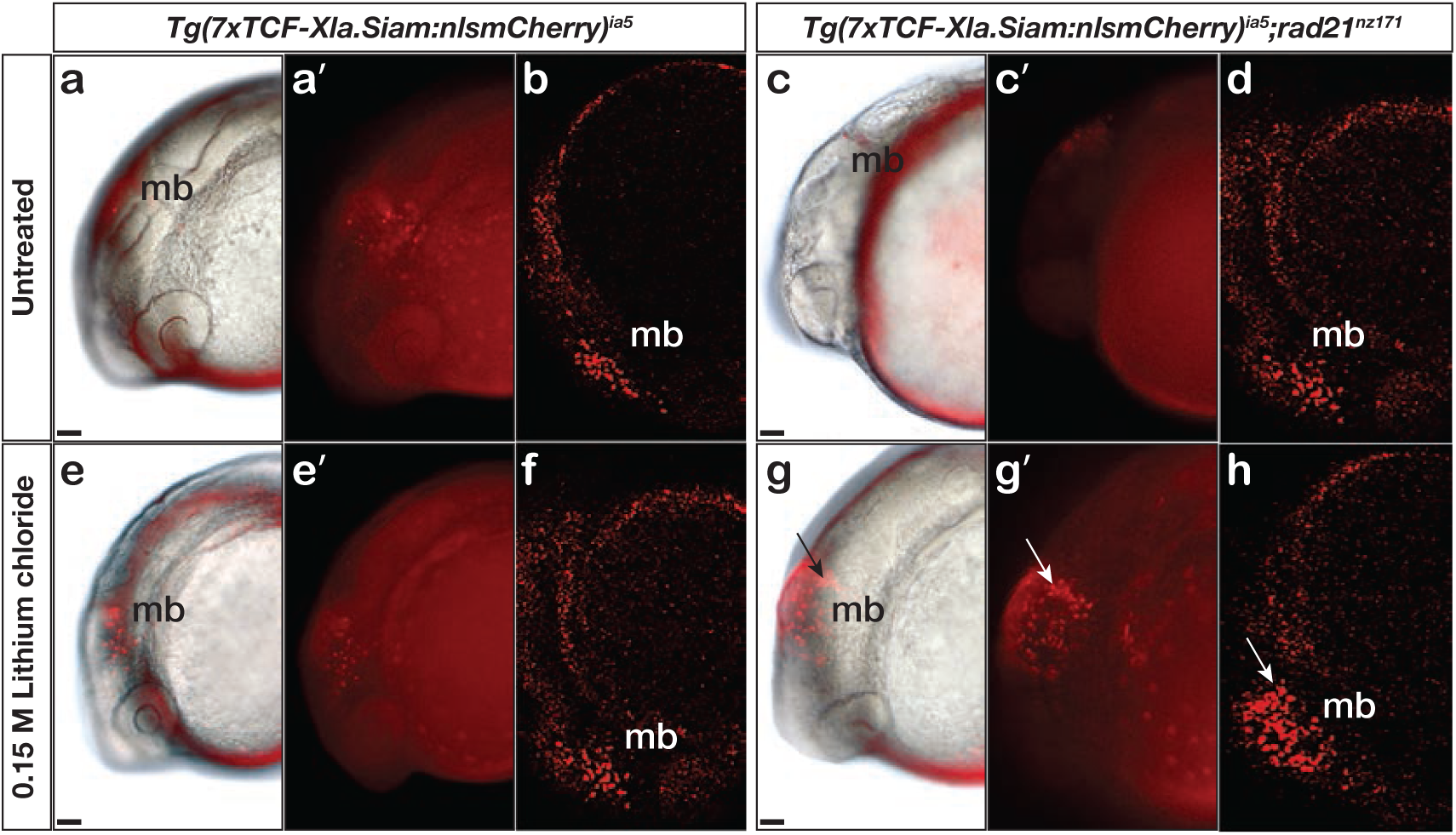
Zebrafish *rad21* cohesin mutants show increased sensitivity to Wnt stimulation. Wnt reporter *Tg(7xTCF-Xla*.*Siam:nlsmCherry)*^*ia5*^ control embryos and *Tg(7xTCF-Xla*.*Siam:nlsmCherry)*^*ia5*^;*rad21*^nz171^ cohesin mutant embryos were treated with 0.15 M LiCl from 4 hpf to 20 hpf. **a, a’** Brightfield/fluorescent and **b** confocal images of *Tg(7xTCF-Xla*.*Siam:nlsmCherry)*^*ia5*^ transgenic control embryos. **c, c’** Brightfield/fluorescent and **d** confocal *Tg(7xTCF-Xla*.*Siam:nlsmCherry)*^*ia5*^;*rad21*^nz171^ cohesin mutant embryos. Both control and *rad21*^nz171^ genotypes show constitutive Wnt reporter activity in the midbrain at 20 hpf. Control brightfield/fluorescent (**e, e’**) and confocal (**f**) images compared with *rad21*^*nz171*^ brightfield/fluorescent (**g, g’**) and confocal (**h**) images show that in the cohesin mutation background, Wnt reporter activity is enhanced in the midbrain (arrow) upon LiCl treatment, relative to controls. mb, midbrain. Scale bar, 50 µm. For *Tg(7xTCF-Xla*.*Siam:nlsmCherry)*^*ia5*^ control embryos n=15/15; and for *Tg(7xTCF-Xla*.*Siam:nlsmCherry)*^*ia5*^;*rad21*^nz171^ cohesin mutant, n=4/4 with the same expression pattern.

The identification of DNA damage repair inhibitors in our screen agrees with previous studies showing synthetic lethality of PARP inhibition with cohesin mutation [33]. Differential sensitivity of the cohesin deletion cell lines to PI3K/AKT/mTOR inhibitors is consistent with the observed nucleolar deficiencies in cohesin mutant cells (Supplementary Fig. 4). The PI3K/AKT/mTOR pathway stimulates ribosome biogenesis and rDNA transcription; because rDNA is contained in nucleoli, it is likely that rDNA transcription and ribosome production is already compromised in the cohesin gene deletion cell lines. We had previously shown that BET inhibition is effective in blocking precocious gene expression in the chronic myelogenous leukemia cell line K562 containing a *STAG2* mutation [45]. Growth inhibition of cohesin-deficient MCF10A cells by I-BET-762 (Fig. 3f, Supplementary Fig. 6e) reinforces the idea that targeting BET could be therapeutically effective in cohesin-mutant cancers. Interestingly, we found that LY2090314, a GSK3 inhibitor and stimulator of the Wnt signaling pathway, inhibits all three cohesin deletion lines (Table 1, Fig. 3g, Supplementary Fig. 6f). LY2090314 also inhibited the growth of K562 STAG2 R614* mutant leukemia cells that we had previously characterized [45] (Supplementary Fig. 6g). Wnt signaling appears to act upstream of cohesin [46, 47], and also to be primarily downregulated downstream of cohesin mutation [48-50]. Therefore, we were prompted to further investigate why Wnt stimulation via GSK3 inhibition might cause lethality in cohesin-deficient cells.

### Stabilization of β-catenin in cohesin-deficient MCF10A cells

In “off” state of canonical Wnt signaling, β-catenin forms a complex including Axin, APC, GSK3, and CK1 proteins. β-catenin phosphorylated by CK1 and GSK3 is recognized by the E3 ubiquitin ligase β-Trcp, which targets β-catenin for proteasomal degradation. Activation of Wnt signaling by ligand binding, or by GSK3 inhibition, releases β-catenin, allowing it to accumulate in the nucleus where it binds TCF to activate Wnt target gene transcription [51, 52]. We found that there was no difference in GSK3 levels between parental MCF10A cells and the cohesin-deficient clones, and upon treatment of cells with LY2090314, levels of GSK3 were reduced in all cells as expected (Supplementary Fig. 7). In contrast, in untreated cells we found that β-catenin is stabilized in all three cohesin-deficient clones compared with parental MCF10A cells. When Wnt signaling is in the “off” state, phospho-Ser33/37/Thr41 marks β-catenin targeted for degradation. Membrane-associated phospho-Ser33/37/Thr41-β-catenin was increased in cohesin-deficient clones, and disappeared upon treatment with LY2090314 (Fig. 4a). Phospho-Ser675 β-catenin, the active form of β-catenin that is induced upon Wnt signaling, was noticeably increased in the cytoplasm of cohesin-deficient cells following treatment with LY2090314 (Fig. 4b, c). The results imply that inactive β-catenin that would normally be targeted for degradation is instead stabilized in cohesin-deficient cells, and is available to be converted into the active form following inhibition of GSK3.

Immunofluorescence labeling of cells showed that in LY2090314-treated cells, the stabilized active phospho-Ser675 β-catenin (and total β-catenin, Supplementary Fig. 8a) mainly locates to puncta in the cytoplasm (Fig. 4d). In contrast, no β-catenin accumulation was observed in DMSO-treated cells (Supplementary Fig. 8b). β-catenin-containing puncta were also observed upon WNT3A treatment of cohesin-deficient clones (Supplementary Fig. 8c). This indicates that Wnt stimulation rather than a secondary effect of LY2090314 is responsible for β-catenin accumulation. We could not reliably detect an increase of β-catenin in the nucleus of cohesin-deficient MCF10A clones by immunofluorescence (Fig. 4d), therefore we decided to use transcriptional response to determine the consequences of β-catenin stabilization for Wnt signaling.

### Cohesin-deficient leukemia cells are hypersensitive to Wnt signaling

RNA sequencing analysis of the cohesin gene deletion clones compared with parental MCF10A cells revealed that gene expression changes strongly track with the identity of the deleted cohesin gene (Supplementary Fig. 9a). However, expressed transcripts encoding Wnt signaling pathway components did not cluster differently in cohesin-deficient clones compared with parental MCF10A cells (Supplementary Fig. 9b), and cohesin mutation-based clustering remained dominant. We reasoned that stimulation of Wnt signaling in a more responsive cell type might be necessary to determine how stabilized β-catenin in cohesin-deficient cells affects transcription.

Myeloid leukemias are frequently characterized by cohesin mutations, particularly in *STAG2*. Furthermore, activation of Wnt signaling is associated with transformation in AML [53-55] and AML patients were identified with mutations in *AXIN1* and *APC* that lead to stabilization of β-catenin [56]. CMK is a Down Syndrome-derived megakaryoblastic cell line that typifies myeloid leukemias that are particularly prone to cohesin mutation [57]. We edited CMK to contain the STAG2-AML associated mutation R614* [45] and selected single clones with complete loss of the STAG2 protein to create the cell line CMK-*STAG2-/-* (Supplementary Fig. 10a-c).

Immunofluorescence showed that β-catenin is increased by 20% in the nuclei of CMK*-STAG2-/-* cells relative to parental cells upon WNT3A stimulation (Fig. 5a, b). To determine immediate early transcriptional responses to Wnt signaling, RNA-sequencing was performed on CMK-*STAG2-/-* and parental CMK cells at baseline and after stimulation with WNT3A for 4 hours. Around 76% more genes were upregulated in response to WNT3A in CMK-*STAG2-/-* compared with CMK parental cells (616 in CMK-*STAG2-/-* vs 350 in CMK parental, FDR ≤0.05, Fig. 5c), while around the same number were downregulated. About one quarter of differentially expressed genes overlapped between CMK-*STAG2-/-* and CMK parental cells.

Strikingly, 404 genes that were not Wnt-responsive in CMK parental cells became Wnt-responsive upon introduction of the *STAG2* R614* mutation (Fig. 5d). Genes upregulated in CMK-*STAG2-/-* markedly increased the number of Wnt-sensitive biological pathways following WNT3A treatment compared with parental CMK cells (Fig. 5e). A strongly upregulated cluster of 244 transcripts in CMK-*STAG2-/-* included genes encoding signaling molecules (JAG2, IL6ST, SEM3G) and transcription factors (SP7, KLF3, SMAD3, EP300 and EPHA4) (Supplementary Fig. 10d). Genes in this cluster were enriched for LEF1 and TCF7 binding indicating their potential to be directly regulated by β-catenin. Enriched biological pathways included Wnt signaling, cell cycle and metabolism (Supplementary Fig. 10e). Pathway analyses of genes significantly downregulated only in CMK-*STAG2-/-* also showed enrichment for metabolism (Supplementary Fig. 10f, g).

Overall, our results show that CMK-*STAG2-/-* cells are exquisitely sensitive to Wnt signaling. Introduction of the *STAG2* R614* mutation amplified expression of Wnt-responsive genes and sensitized genes and pathways that are not normally Wnt-responsive in CMK. This sensitivity could be due at least in part to stabilized β-catenin.

### Conservation of WNT sensitivity in cohesin-deficient zebrafish

To determine if enhanced Wnt sensitivity is conserved in a cohesin-deficient animal model, we used our previously described zebrafish mutant, *rad21*^nz171^ [58], which has a nonsense point mutation in the *rad21* gene that eliminates Rad21 protein. To provide a readout for Wnt signaling, we used transgenic zebrafish carrying a TCF/β-catenin reporter in which exogenous Wnt stimulation induces nuclear mCherry: *Tg(7xTCF-Xla*.*Siam:nslmCherry)*^*ia5*^ [59]. The Wnt reporter construct, which drives nuclear mCherry red fluorescence in Wnt-responsive cells, was introduced into zebrafish carrying the *rad21*^nz171^ mutation by crossing. Zebrafish embryos heterozygous for *Tg(7xTCF-Xla*.*Siam:nslmCherry)*^*ia5*^, and either homozygous for *rad21*^nz171^ or wild type, were exposed to 0.15 M LiCl, an agonist of the Wnt signaling pathway (Fig. 6). Expression of mCherry in the midbrain of embryos was detected at 20 hours post-fertilization (hpf) by epifluorescence and confocal imaging. A constitutive low level of mCherry is present in the developing midbrain of both untreated wild type and *rad21*^nz171^ embryos (Fig. 6a-d). LiCl treatment dramatically increased the existing midbrain mCherry expression in *rad21*^nz171^ mutants compared with wild type embryos (Fig. 6e-h).

The results show that Wnt signaling is sensitized in *rad21* cohesin mutant zebrafish embryos, similar to our observations in MCF10A and CMK cell lines. Our findings agree with previous work showing that canonical Wnt signaling is hyperactivated in cohesin loader *nipblb*-loss-of-function zebrafish embryos [60]. Altogether, the results suggest that hyperactivation of Wnt signaling is a conserved feature of cohesin deficient cells, and that enhanced sensitivity to Wnt is at least in part due to stabilization of β-catenin.

## Discussion

The cohesin complex and its regulators are encoded by several different loci, and genetic alterations in any one of them may occur in up to 26% of patients included in The Cancer Genome Atlas (TCGA) studies [61]. Therefore, we were motivated to identify compounds that interfere with cell viability in more than just one type of cohesin mutant. The generation of *RAD21, SMC3* and *STAG2* cohesin mutations in the breast epithelial cell line MCF10A resulted in mild cell cycle and nucleolar phenotypes that are consistent with the many cellular roles of cohesin. Differences between *RAD21* and *SMC3* heterozygotes vs *STAG2* homozygotes could be explained by the fact that RAD21 and SMC3 are obligate members of the cohesin ring, whereas STAG2 can be compensated by STAG1.

Synthetic lethal sensitivities of cohesin mutant cells that emerged from our screen mostly related to the phenotypes of cohesin mutant cells, and/or their previously identified vulnerabilities. For example, cohesin mutant cells were sensitive to inhibitors of the PI3K/Akt/mTOR pathway that feeds into ribosome biogenesis, epigenetic inhibitors that could interfere with cohesin’s genome organization and gene expression roles, and a limited sensitivity to PARP inhibitors. The observed sensitivity to PI3K/AKT/mTOR inhibitors is consistent with the nucleolar disruption present in cohesin-deficient cell lines, and with cohesin’s known involvement in rDNA transcription and ribosome biogenesis [38]. We have also previously described evidence for sensitivity of cohesin-deficient cell lines to bromodomain (BET) inhibition [45]. However, the Wnt agonist LY2090314, which mimics Wnt activation by inhibiting GSK3-β [62], emerged as a novel class of compound that inhibited the growth of all three cohesin mutants tested. Because there is pathway convergence between Wnt signaling and the PI3K/AKT/mTOR pathway [63] it is possible that these pathways represent a common avenue of sensitivity. The identification of PI3K/AKT/mTOR inhibitors in the screen supports this notion.

Wnt signals are important for stem cell maintenance and renewal in multiple mammalian tissues [52]. Canonical Wnt signaling is required for self-renewal of leukemia stem cells (LSCs) [55], and is reactivated in more differentiated granulocyte-macrophage progenitors (GMPs) when they give rise to LSCs [53]. Wnt signaling is an important regulator of hematopoietic stem cell (HSC) self-renewal as well [64]. Our results suggest that cohesin deficient cells are sensitive to Wnt agonism because they already have stabilization of β-catenin. Interestingly, multiple experimental systems showed that dose-dependent reduction in cohesin function causes HSC expansion accompanied by a block in differentiation [65, 66]. It is possible that the effects of cohesin deficiency on HSC development could be in part due to enhanced Wnt signaling.

Cohesin mutations are particularly frequent (53%) in Down Syndrome-associated Acute Megakaryoblastic Leukemia (DS-AMKL) patients [57]. In the edited DS-AMKL cell line *STAG2*-CMK, there was a dramatically enhanced immediate early transcriptional response to WNT3A, including induction of hundreds of genes that are not normally responsive to WNT3A in this cell line. It is possible that STAG2 deficiency in CMK leads to an altered chromatin structure that sensitizes genes to Wnt signaling. However, we observed increased nuclear localization of β-catenin in *STAG2*-CMK, increasing the likelihood that stabilization of β-catenin in *STAG2*-CMK cells accounts for amplified gene expression in response to WNT3A. Interestingly, DS-AMKL leukemias are often associated with amplified Wnt signaling caused by Trisomy 21 [67] suggesting a possible synergy between cohesin mutations and Wnt signaling dysregulation in DS-AMKL.

We observed an enhanced Wnt reporter response to Wnt activation in *rad21* mutant zebrafish, arguing that Wnt sensitivity upon cohesin mutation is a conserved phenomenon. Previous research shows that cohesin genes are at once both targets [46, 68] and upstream regulators [47, 49, 50, 60] of Wnt signaling. For example, depletion of cohesin loader *nipbl* in zebrafish embryos was reported to downregulate Wnt signaling at 24 hpf [69], but to upregulate it at 48 hpf [60]. What determines the directionality of cohesin’s effect on Wnt signaling is unclear. It is possible that feedback loops operate differently in cell-and signal-dependent contexts [70] in cohesin-deficient backgrounds.

A remaining question is exactly what causes β-catenin stabilization and Wnt hyperactivation in cohesin mutant cells. One possibility is that stabilization of β-catenin upon cohesin deficiency is linked to energy metabolism. For example, in developing amniote embryos, glycolysis in actively growing cells of the embryonic tail bud raises the intracellular pH, which in turn causes β-catenin stabilization [71, 72]. This mode of glycolysis is similar to that observed in cancer cells and is known as the Warburg effect [73, 74]. Metabolic and oxidative stress disturbances in cohesin mutant cells observed in this study and others [38, 75] might similarly influence β-catenin stability and in turn, sensitivity to incoming Wnt signals. Further research will be needed to determine how β-catenin is stabilized in cohesin deficient cells, whether β-catenin stabilization is part of a transition to malignancy, and if the associated Wnt sensitivity represents a therapeutic opportunity in cohesin mutant cancers.

## Supporting information

Active compounds from primary screen

Active compounds from secondary screen

## Material and Methods

### Cell culture

MCF10A and its cohesin-deficient derivatives were maintained in Dulbecco’s modified Eagle medium (DMEM) (Life Technologies) supplemented with 5% horse serum (Life Technologies), 20 ng/mL of epidermal growth factor (EGF) (Peprotech), 0.5 mg/mL of hydrocortisone, 100 ng/mL of cholera toxin and 10 μg/mL of insulin (local pharmacy). All supplements were purchased from Sigma-Aldrich unless otherwise stated. K562 cells were maintained in Isocove’s Modified Dulbecco’s Media (IMDM) (Life Technologies) containing 10% fetal bovine. CMK cells were maintained in RPMI 1640 media containing 10% fetal bovine serum. CMK cells and adherent K562-*STAG2* null line were detached for subculture using trypsin-EDTA (0.005% final concentration, Life Technologies). All cells were cultured at 37 °C at 5% CO2.

### Generation of isogenic cell lines using CRISPR-CAS9 editing

We used CRISPR-CAS9 sgRNAs targeting the 5’ and 3’ UTR regions of *RAD21, SMC3* and *STAG2* gene to create MCF10A *RAD21+/-*, MCF10A *SMC3+/-*, and MCF10A *STAG2-/-*. Specific sgRNAs were cloned into the px458 plasmid [76] and transfected into MCF10A cells. Single GFP-positive cells were isolated into 96-well plates using a FACSAriaII (Becton Dickinson) and clonally expanded. PCR screening and Sanger sequencing identified cells with heterozygous deletion of *RAD21* or *SMC3* and homozygous deletion of *STAG2*. Primer and guide sequences are provided in Supplementary Tables 3 and 4. Editing of K562 to create STAG2 R614* mutation has been described previously [45]. The CMK line with the STAG2 R614* mutation was generated using the same sgRNA and method.

### Quantitative PCR (RT-qPCR)

Total RNA was extracted using NucleoSpin RNA kit (Machery-Nagel). cDNA was synthesized using qScript™ cDNA SuperMix (Quanta Biosciences). RT-qPCR was performed on a LightCycler® 480 II (Roche Life Science) using SYBR® Premix Ex Taq(tm) (Takara). Expression values relative to reference genes *cyclophilin* and *glyceraldehyde 3-phosphate dehydrogenase* (*GAPDH*) were derived using qBase Plus (Biogazelle). Primer sequences are provided in Supplementary Table 5.

### Antibodies

Primary antibodies used are as follows: anti-RAD21 (ab992), anti-SMC3 (#5696, CST), anti-STAG2 (ab4463), anti-γ-Tubulin (T5326; Sigma-Aldrich) in 1:5000, anti-fibrillarin (ab5821), anti-nucleolin (ab13541), anti-gamma H2AX (ab26350), anti-TP53 (ab131442), anti-β-catenin (#9562, CST), anti-phospho-β-catenin (Ser675) (#9567, CST), anti-phospho-phospho-β-catenin (Ser33/37/Thr41) (#9561, CST). Secondary antibodies used for immunofluorescence were anti-mouse Alexa Fluor 488 (1:2000, Life Technologies), anti-rabbit Alexa Fluor 568 (1:2000, Life Technologies). All antibodies were used in 1:1000 for immunoblotting or immunofluorescence unless otherwise stated.

### Immunoblotting

Immunoblotting was carried out as described previously [77, 78]. Primary antibodies were detected using IRDye 800CW Donkey anti-Goat IgG and IRDye 680CW Goat anti-mouse IgG, IRDye 800CW Goat anti-rabbit IgG (LICOR). LI-COR Odyssey® and LI-COR Image Studio software was used to image and quantify blots.

### Proliferation assays

MCF10A parental and isogenic cohesin deficient cell lines were seeded in 96-well plates at 2,000 cells per well. Cell confluence was monitored by time-lapse microscopy using IncuCyte™ FLR with a 10X objective for 5 days.

### Cell cycle analysis

Cells were synchronized using double thymidine block as described previously [78]. Samples were harvested, fixed and stained with 10 μg/ml propidium iodide (PI) and 250 μg/ml RNase A, 37 °C for 30 min. Cells were then analyzed using a Beckman Coulter Gallios Flow Cytometer.

### Immunofluorescence and imaging

Cells stained for FBL, gH2AX, and TP53 were imaged using an Opera Phenix high content screening system, with 63x water objectives in confocal mode. Spot enumeration and signal intensities were analyzed using Harmony software (PerkinElmer). CMK cells were fixed with 4% (v/v) paraformaldehyde in PBS for 10 minutes, washed, stained with Hoechst 33342 (1 μg/mL) for 10 min. CMK cells were spun onto slides using SHANDON cytospin prior to fixation. Cells were blocked with 2% (w/v) bovine serum albumin in PBS, then incubated with primary antibody overnight at 4 °C. The next day, cells were washed with PBS and incubated with secondary antibody at room temperature for 1 hour. Cells were then washed and mounted with microscope slides using ProLong Gold Anti-fade (Life Technology) or DAKO mounting medium. Imaging was done using a Nikon C2 confocal microscope with NIS Elements and images were processed and quantified using Image J or FIJI software.

### High-throughput compound screen

3,009 compounds from the FDA-approved, kinase inhibitors and epigenetic libraries (Compounds Australia, Griffith University, Australia) were screened against MCF10A parental and MCF10A cohesin-deficient clones. MCF10A parental cells were seeded at 600 cells/well, *RAD21+/-* at 800 cells/well, *SMC3+/-* at 1,200 cells/well and *STAG2-/-* at 700 cells/well, into 384-well CellCarrier-384 Ultra microplates (PerkinElmer) with EL406 microplate washer dispenser (BioTek). 24 hours later, growth media was aspirated from the plates and 35uL of fresh MCF10A complete growth media was added into each well using BioTek EL406TM Microplate washer dispenser. 5uL of 8x concentrated compound solution (diluted in MCF10A complete growth media) was added into each well with a JANUS automated liquid handling system (PerkinElmer). At the time of drug treatment, one untreated plate was retained to determine the cell number at t=0. For the rest of the assay plate, compounds were added in 40 µL of complete medium per well using an Echo 550 Liquid Handler (Labcyte). Positive (Campthothecin, 40 nM) and negative controls (DMSO 0.1%) were added to each assay plate for quality control. Duplicated control plates of each cell line were also prepared for cell count quantification prior to the addition of drugs, to be used in the growth rate inhibition (GR) metrics. After 48 hours, control and treated cells in were fixed and stained simultaneously using 4% PFA/ 0.5ug/mL DAPI/ 0.1% Triton X-100 solution. Cells were imaged using an Opera Phenix high content screening system and analyzed using Columbus software (PerkinElmer) using 20x water objectives. To determine compound activity, dose-response curves for each compound was plotted and using an R package, GRmetrics [79]. Synthetic lethal candidate compounds were selected and ranked based on the differential area over curve (AOC) metrics [44] derived from dose-response curves of MCF10A parental and cohesin deficient cell lines. Compounds that caused ≥30% growth inhibition (AOC ≥ 0.15) in cohesin deficient clones compared with parental MCF10A cells were selected as hits. A secondary screen was performed with 85 candidate hits identified from primary screen. Activity of the selected compounds were tested in eleven concentration (0.5 nM to 10 µM). Secondary screen hit compounds were selected based on the same threshold used in primary screen.

### Cell viability assays to validate compounds identified from the screen

Individual compounds were purchased from Sigma-Aldrich, Selleck Chem or MedChemExpress and dissolved in DMSO at recommended concentrations. MCF10A parental cells were seeded in 96-well plates at 3,000 cells/well, *RAD21+/-* at 4,500 cells/well, *SMC3+/-* at 5,000 cells/well and *STAG2-/-* at 3,500 cells/well. Cells were incubated for 24 hours then treated with compounds for 48 hours. DAPI-stained cells were counted using a Lionheart FX automated microscope (BioTek). K562 parental and *STAG2*-/-cells were seeded at 2,000 cells per well in 96 well plates, incubated with the compound for 48 hours after which viability was measured using 3-(4,5-dimethylthiazol-2-yl)-2,5-diphenyltetrazolium bromide or MTT.

### RNA sequencing and analyses

Total RNA was extracted using NucleoSpin RNA kit (MACHEREY-NAGEL). MCF10A libraries from three biological replicates of each cell type were prepared using NEBNext® Ultra(tm) RNA Library Prep Kit (Illumina®) and sequenced on HiSeq X by Annoroad Gene Technology Ltd. (Beijing, China), contracted through Custom Science (NZ). CMK lines were treated with 200 ng/ml of recombinant human WNT3A (R&D systems) for 4 hours. Libraries were prepared from baseline (non-treated) and WNT3A treated CMK cell lines using Illumina TruSeq stranded mRNA library and sequenced on the HiSeq 2500 V4 at the Otago Genomics Facility (Dunedin, New Zealand). RNA sequencing reads were first trimmed for sequencing adapters and low quality (q<20). Cleaned reads were then aligned to the human genome GRCh37 (hg19) using HISAT2 version 2.0.5 with gene annotation from Ensembl version 75. Read counts were retrieved by exon and summarized by gene using featureCount [80] version v1.5.3. Differentially expressed genes in the *STAG2-/-* mutants versus wild type were identified using DESeq2 [81]. P-values were adjusted for multi-test following Benjamini-Hochberg methology. Pathways analyses were performed using Molecular Signature Databases ReactomePA [82] and clusterProfiler [83] on differentially expressed genes.

### Zebrafish methods and imaging

Wild type (WIK), *Tg(7xTCF-Xla*.*Siam:nslmCherry)*^*ia5*^ [59] and *rad21*^nz171^ [58] mutant fish lines were maintained according to established husbandry methods [84]. Zebrafish embryos were incubated with 0.15 M lithium chloride (LiCl) from 4 hpf for 16 hours. LiCl salt was directly dissolved in embryo water and added to embryos in 6 well plates. 50 embryos were used per treatment group. At 20 hpf, embryos were washed and mounted in 3% methyl cellulose for imaging. Z-stacks were acquired using a Nikon C2 confocal microscope.

### Statistical analyses

Unless otherwise stated, statistical analyses were carried out by a two-tailed Student’s *t* test. A *p* value ≤ 0.05 is considered statistically significant. All statistical analyses were carried out using R and Prism version 7 software (GraphPad).

## Data availability

RNA sequencing data has been deposited at the GEO database under GSE154086.

## Acknowledgements

This work was funded by Health Research Council of NZ awards 15/229 and 19/415 to J.A.H., and a Maurice Wilkins Center for Biodiscovery award to J.A. and J.A.H.

## Author contributions

CVC, JA, SK, GG, PG, AJG, RDH, JAH designed experiments. CVC, JA, SK, GG, KMP, JH performed experiments. CVC, JA, SK, GG, KMP, JH, AJG, AB, PG, RDH, JAH analyzed data. CVC, JA and JAH wrote the paper with input from the other authors. All authors read and approved the final manuscript.

## Supplementary Figure Legends

**Supplementary Figure 1.**
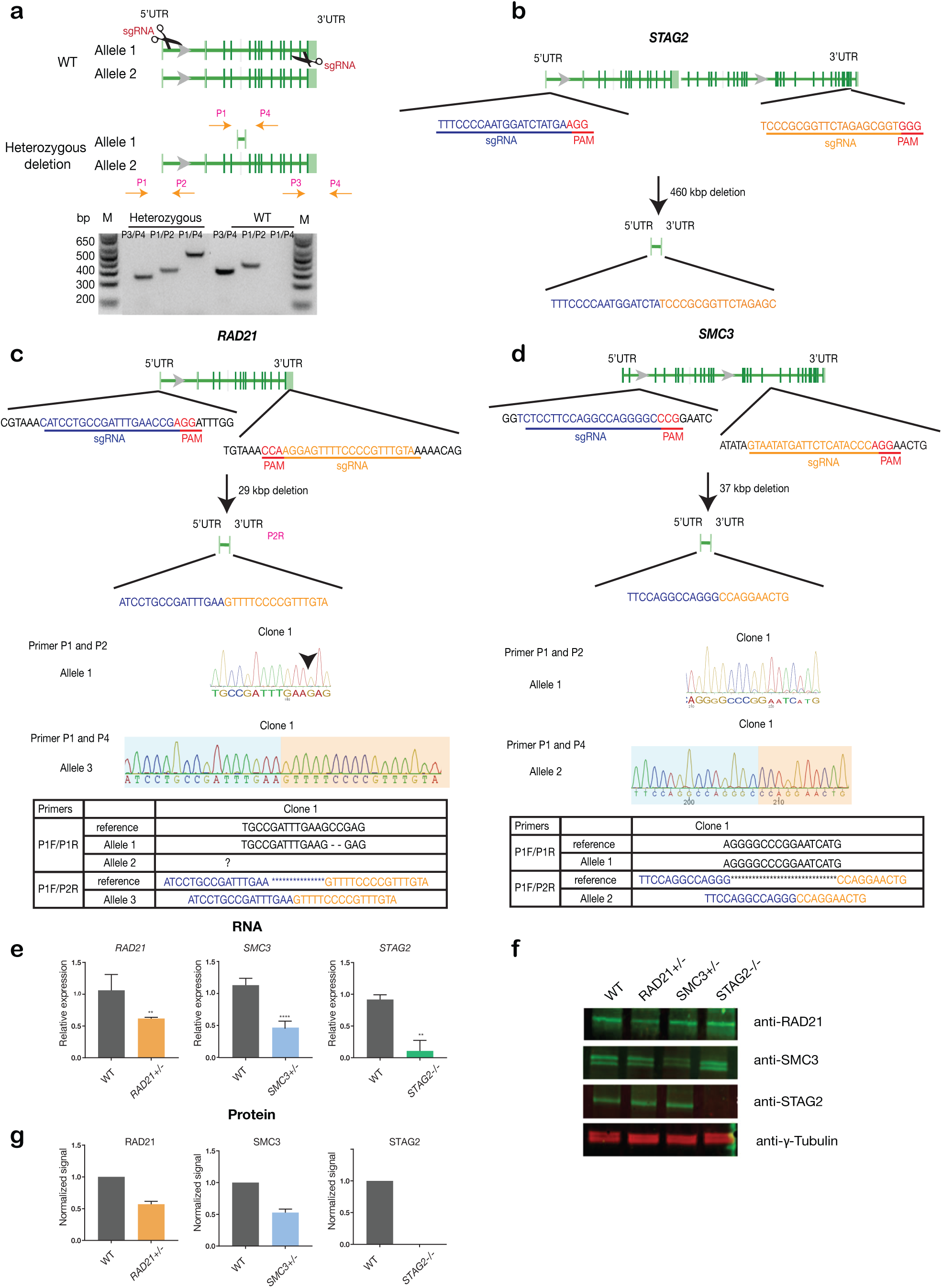
Creation of MCF10A isogenic cell lines with cohesin gene deletions. **a** Top, schematic diagram shows the deletion strategy for genes encoding cohesin subunits RAD21, SMC3 and STAG2 using two sgRNAs targeting the 5’UTR and the 3’UTR of each gene. Bottom, heterozygous clones were identified by PCR using specific primer pairs flanking the deletion region. Representative DNA gel shows the PCR products yielded using specific primer pairs for MCF10A parental and RAD21+/-deletion clone. M, ladder marker. **b-d** Schematic deletion strategy and summary of the allele sequences for the STAG2 homozygous clone, and the RAD21 and SMC3 heterozygous clones. **e** RNA levels of the targeted genes in MCF10A cohesin-deficient clones. f Representative immunoblot and g quantification of cohesin protein levels. γ-tubulin was used as loading control (n = 3 independent experiments, mean ± s.d., **p ≤ 0.01; ****p ≤ 0.0001).

**Supplementary Figure 2.**
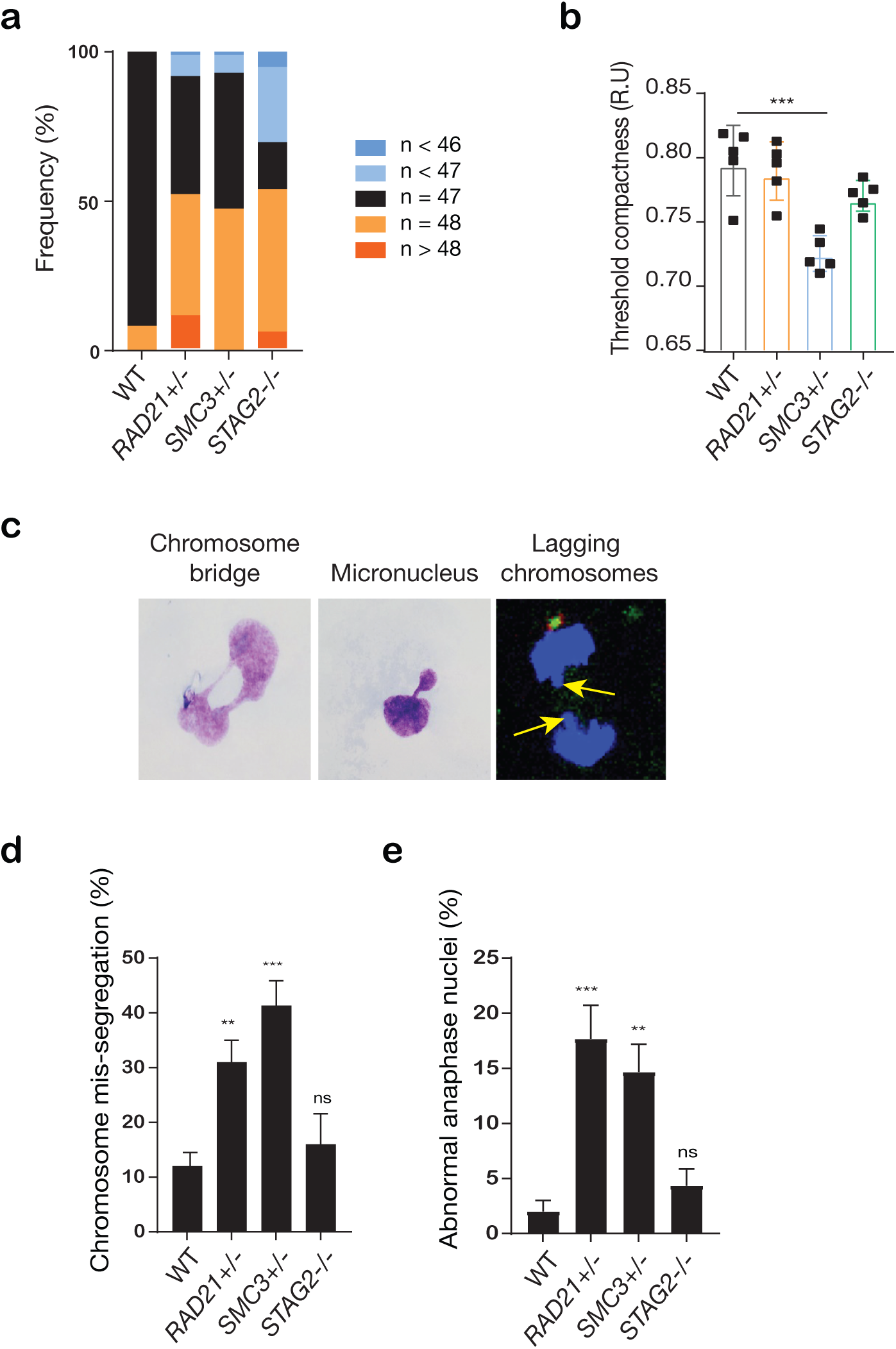
Cohesin deficiency causes a modest increase in chromosome mis-segregation events in MCF10A cells. **a** Frequency of aneuploidy in cohesin-deficient MCF10A clones compared to wild type (WT). **b** Threshold compactness in the nuclei of cohesin-deficient clones compared to wild type MCF10A (R.U. = relative units). **c** Representative images of abnormal interphase nuclei showing chromosome bridges, micronuclei, and lagging chromosomes at anaphase. **d** Quantification of chromosome bridges and micronuclei events in MCF10A cohesin deficient cells. **e** Quantification of chromosome mis-segregation observed in MCF10A cohesin deficient clones. At least 100 mitotic cells were examined. (n = 3 independent experiments, mean ± s.d., **p ≤ 0.01; ***p ≤ 0.0005; ns, not significant).

**Supplementary Figure 3.**
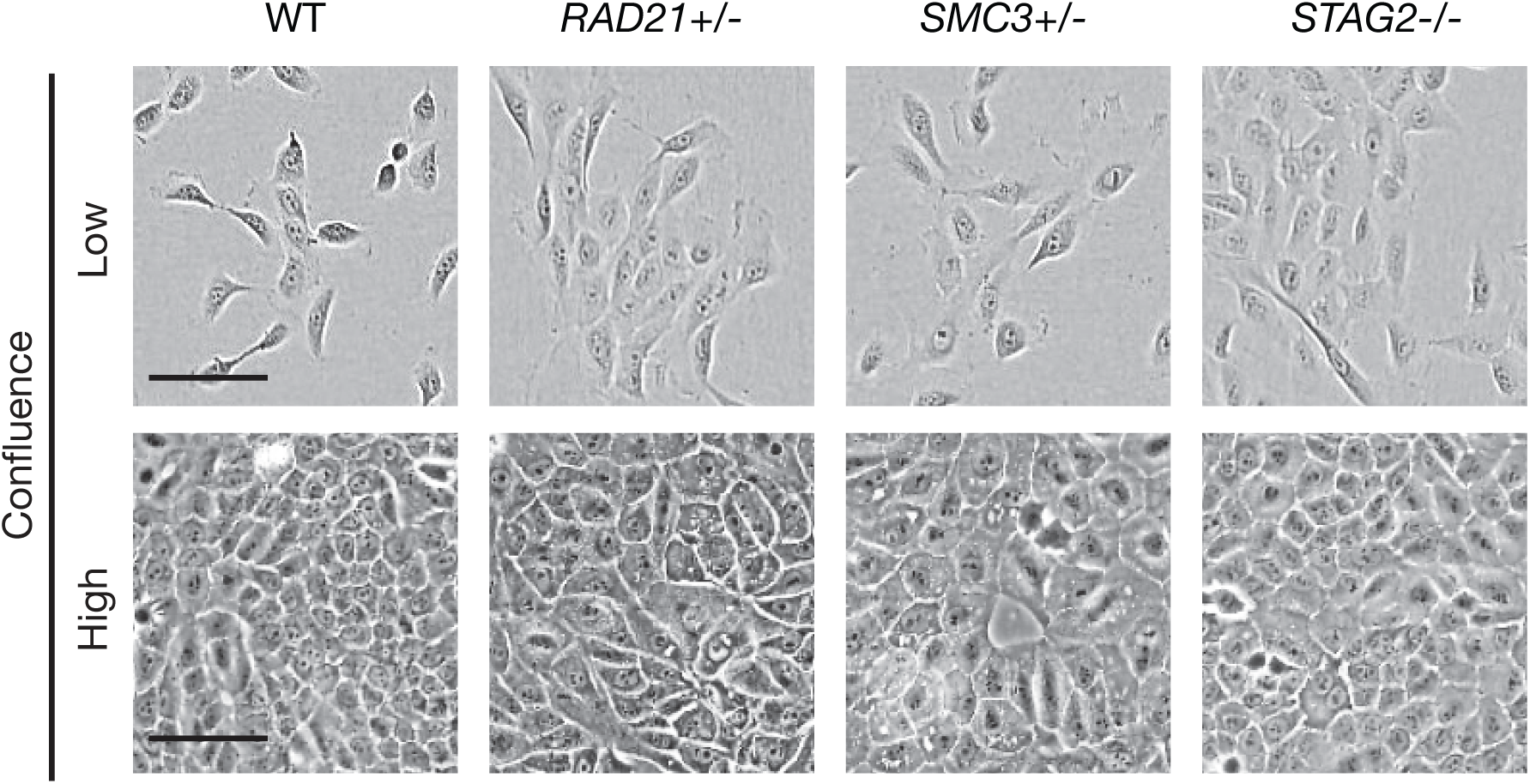
Cell morphology of MCF10A parental and cohesin-deficient cells grown at low and high confluence. MCF10A cells were grown in low (1000 cells/well) and high density (4000 cells/well) in 96-well plates. Cohesin-deficient cells display a cobblestone-like morphology which is similar to MCF10A parental cells. Scale bar, 250 μM.

**Supplementary Figure 4.**
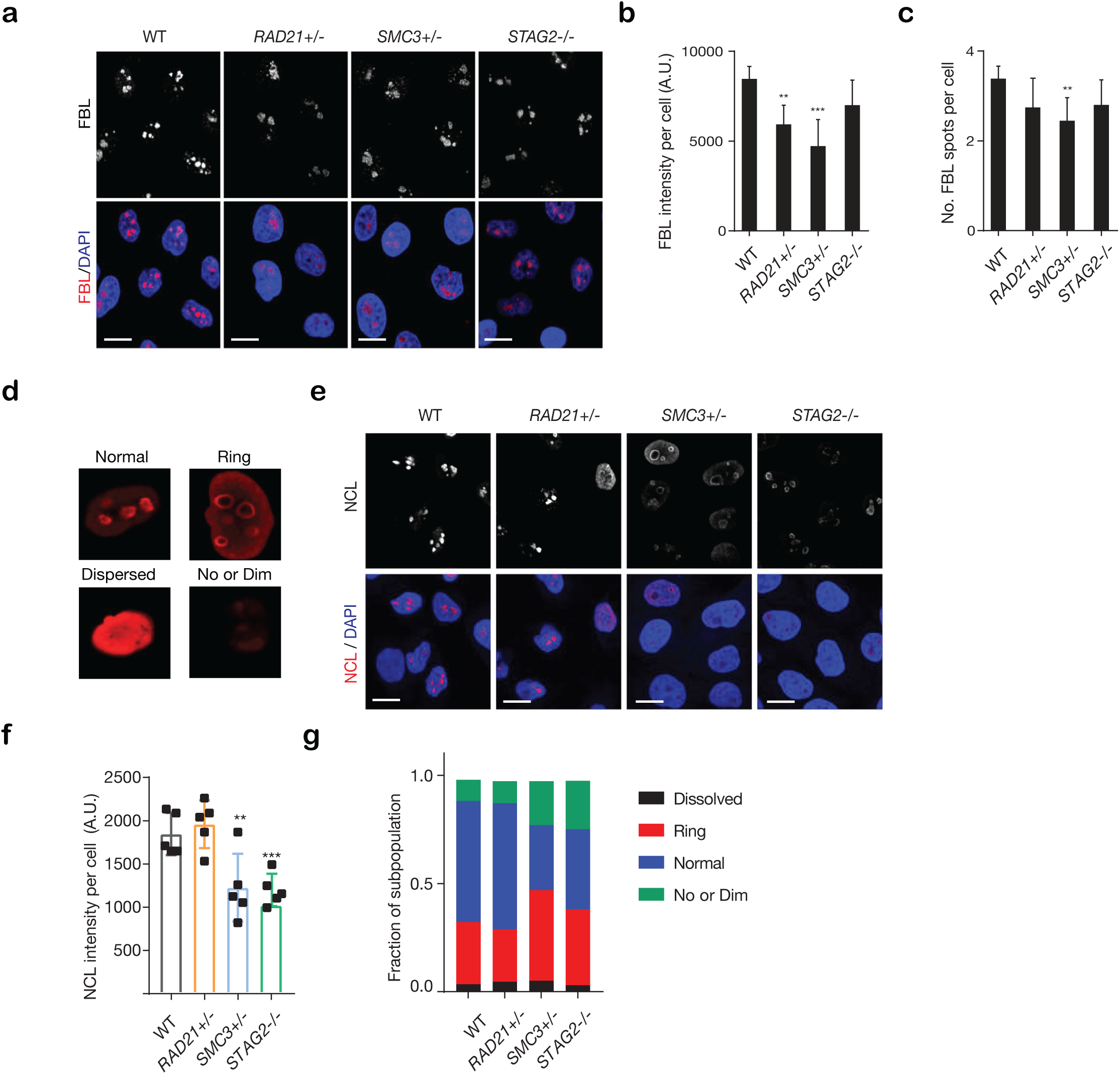
Cohesin deficient cells show altered nucleolar morphology. **a** Representative images of nucleolar morphology as determined using fibrillarin (FBL) as a marker of nucleoli. **b, c** Quantification of FBL intensity and number of FBL spots per cell in each genotype. A minimum of 1000 cells was examined per individual experiment. (n = 3 independent experiments, mean ± s.d., *p ≤ 0.05; **p ≤ 0.01; ***p ≤ 0.0005). **d** Types of nucleolar morphology observed in MCF10A parental cells. **e** Representative images of nucleolar morphology determined by using nucleolin (NCL) as a marker. **f** Quantification of NCL intensity per cell in each genotype. At least 500 cells were examined per individual experiment. n = 3 independent experiments, mean ± s.d., **p ≤ 0.01; ***p ≤ 0.0005). **g** Quantification of NCL morphology classes present in cohesin deficient cells. A.U., Arbitrary unit.

**Supplementary Figure 5.**
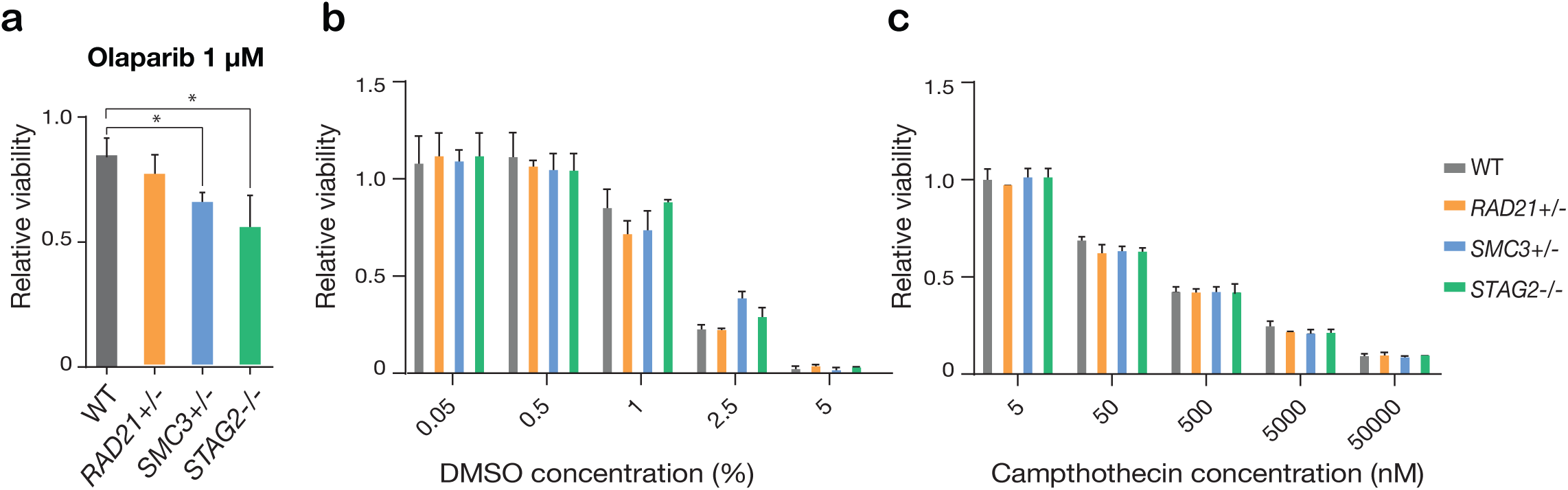
Synthetic lethal screen controls. **a** Cohesin-deficient MCF10A cells show differential sensitivity to the PARP inhibitor, Olaparib, after 48 hours of treatment. **b** DMSO (negative control) and **c** Camptothecin (positive control) titrations were used to establish a cytotoxicity range in MCF10A cells and cohesin-deleted derivatives (n = 2 independent experiments, mean ± s.d.).

**Supplementary Figure 6.**
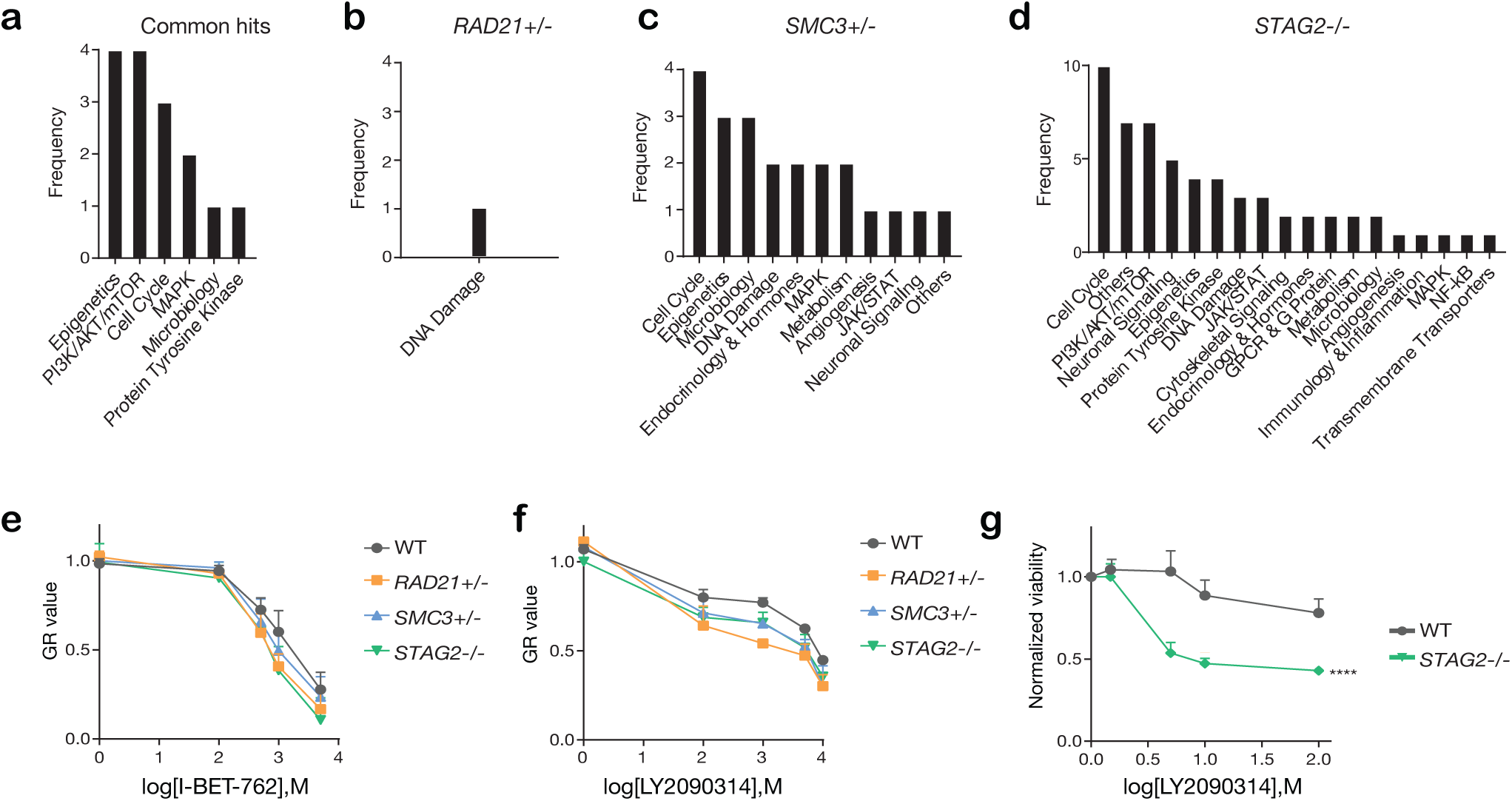
Categories of compounds that differentially inhibit cohesin deficient cells. **a** Common and **b, c, d** unique categories of compounds identified in the primary screen. **e, f** Dose-response curves were generated for validation of I-BET-762 and LY2090314 in MCF10A cells. **g** Dose-response curve of LY2090314 in the K562 STAG2R614* mutant cell line.

**Supplementary Figure 7.**
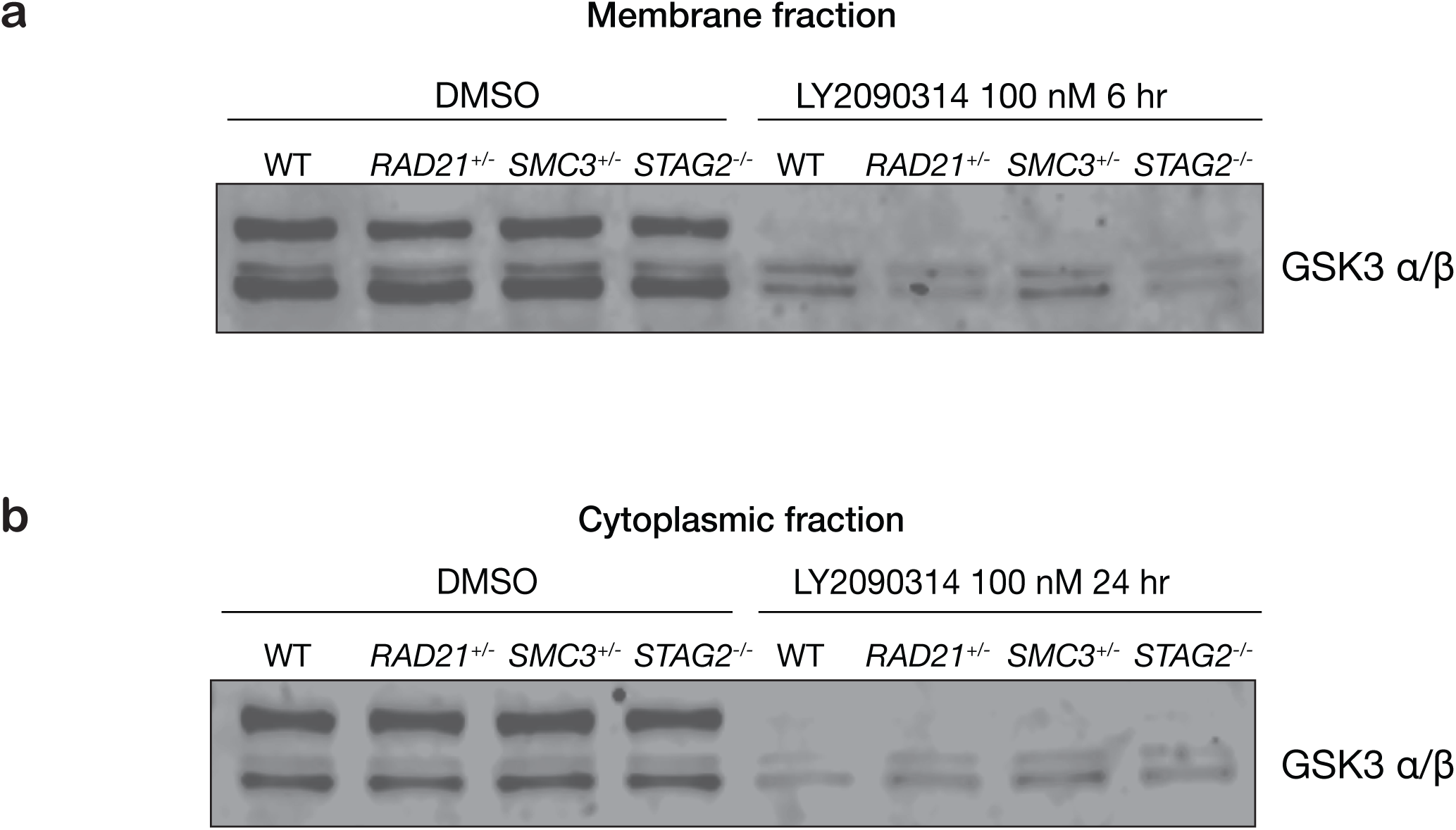
GSK3 levels are unaffected in cohesin-deficient MCF10A cells. Anti-GSK3 immunoblot of membrane fraction from cohesin-deficient cells treated with DMSO or 100 nM LY2090314. **a** Membrane fraction with DMSO and after 6 h LY2090314 treatment. **b** Cytoplasmic fraction with DMSO and after 6h LY2090314 treatment.

**Supplementary Figure 8.**
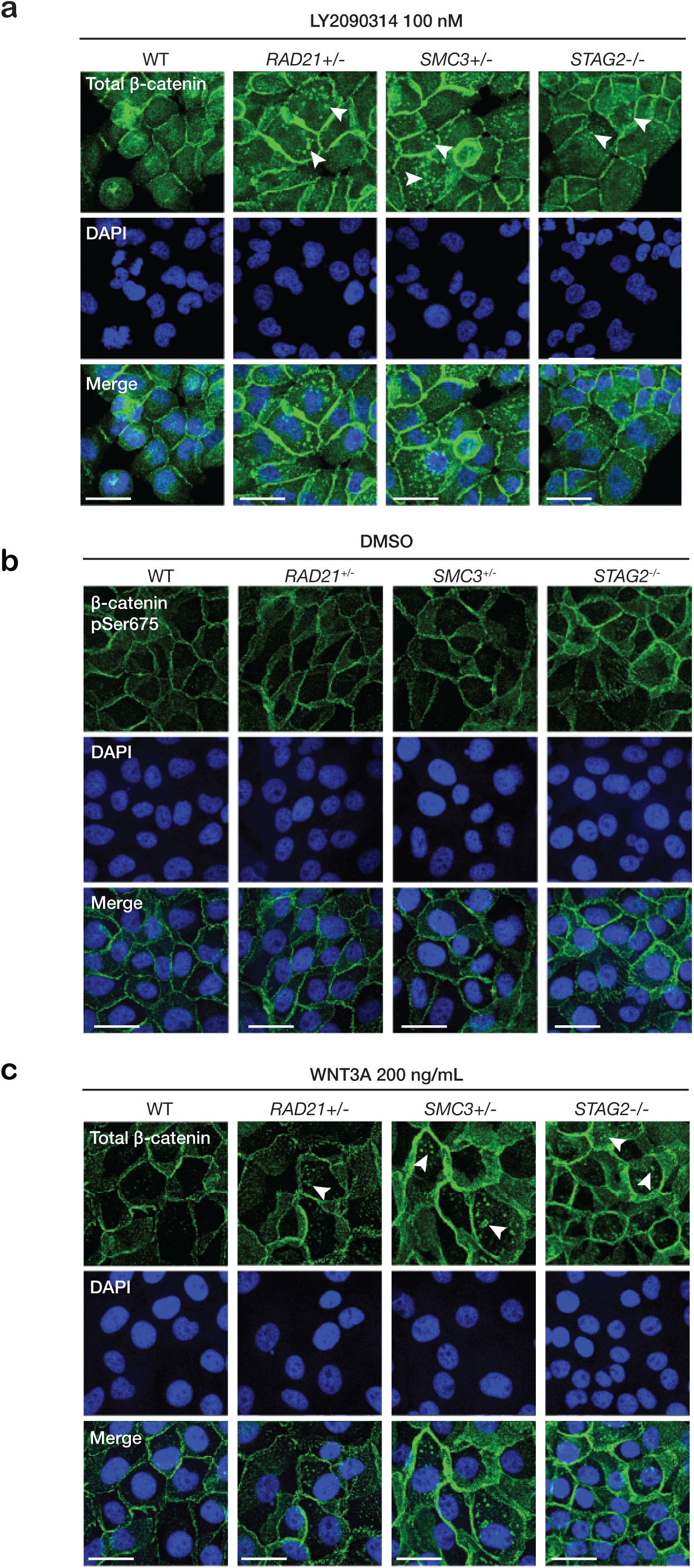
WNT3A phenocopies LY2090314-mediated β-catenin accumulation in the cytoplasm. Immunofluorescence images show that while cell in 0.5% DMSO **(b)** have no β-catenin puncta, treatment with both **a** LY2090314 and **c** WNT3A leads to formation of β-catenin puncta in cytoplasm (arrows). Scale bar = 25 µm.

**Supplementary Figure 9.**
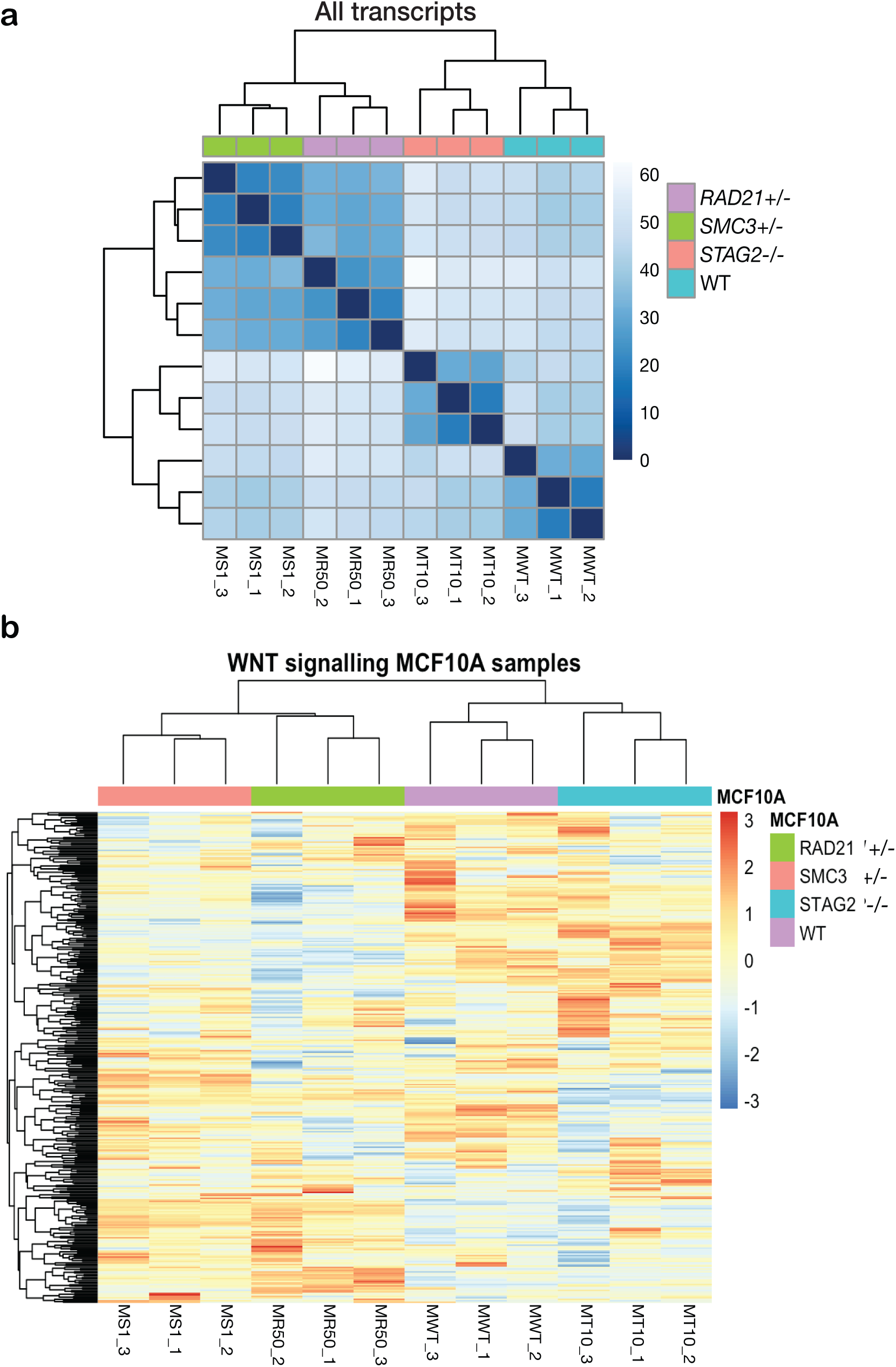
RNA sequencing profiling of cohesin-deficient MCF10A cells. **a** Whole genome correlation matrix of MCF10A cells. The dendrogram shows a common clustering of the biological replicates. There are two main clusters. The STAG2-/-(MT10_1-3) transcriptomes are more similar to the WT (MWT_1-3). SMC3+/-(MS_1-3) and RAD21+/-(MR50_1-3) populate the second cluster. **b** MCF10A gene expression profile of genes involved in WNT signaling pathways according to Gene Ontology annotation (GO:0016055). The heatmap shows the clustering of the scaled (z-score) normalized count (log2RPKM) for each replicate.

**Supplementary Figure 10.**
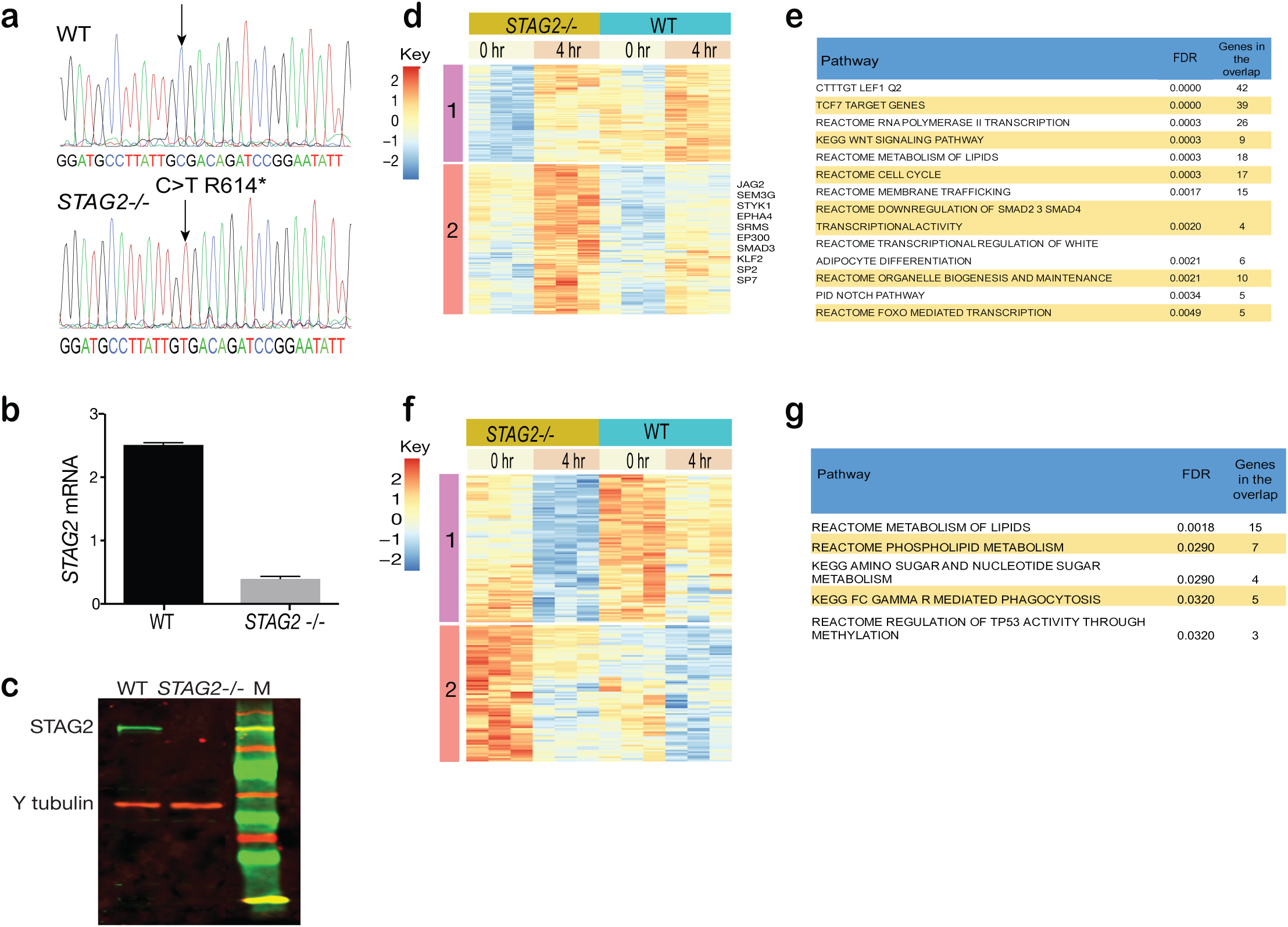
Enhanced sensitivity of cohesin-mutant CMK cells to Wnt stimulation. **a** Sanger sequencing shows successful CRISPR-CAS9 editing introducing the STAG2 R614* C>T mutation site into CMK cells. Reduced STAG2 mRNA **(b)** and protein **(c)** in STAG2 mutant CMK cells. **d** Heatmap showing log2 RPKM values of 404 genes upregulated only in STAG2 mutant cells (FDR ≤ 0.05) following 4 hours of WNT3A treatment. **e** GSEA analyses of the 404 upregulated genes. **f** Heatmap showing log2 RPKM values of 171 genes downregulated only in STAG2 mutant cells (FDR ≤ 0.05) following 4 hours of WNT3A treatment. **g** GSEA analyses of the 171 genes significantly downregulated only in the STAG2 mutant line.

**Supplementary Table 3.**
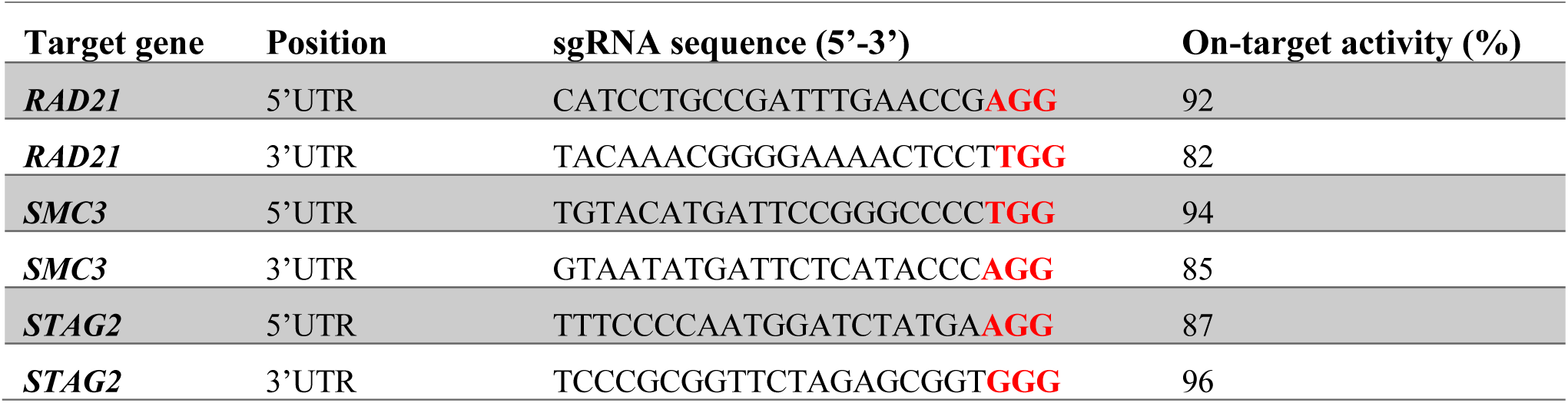
sgRNA sequences targeting each of the cohesin subunit genes. The PAM sequences are highlighted in red.

**Supplementary Table 4.**
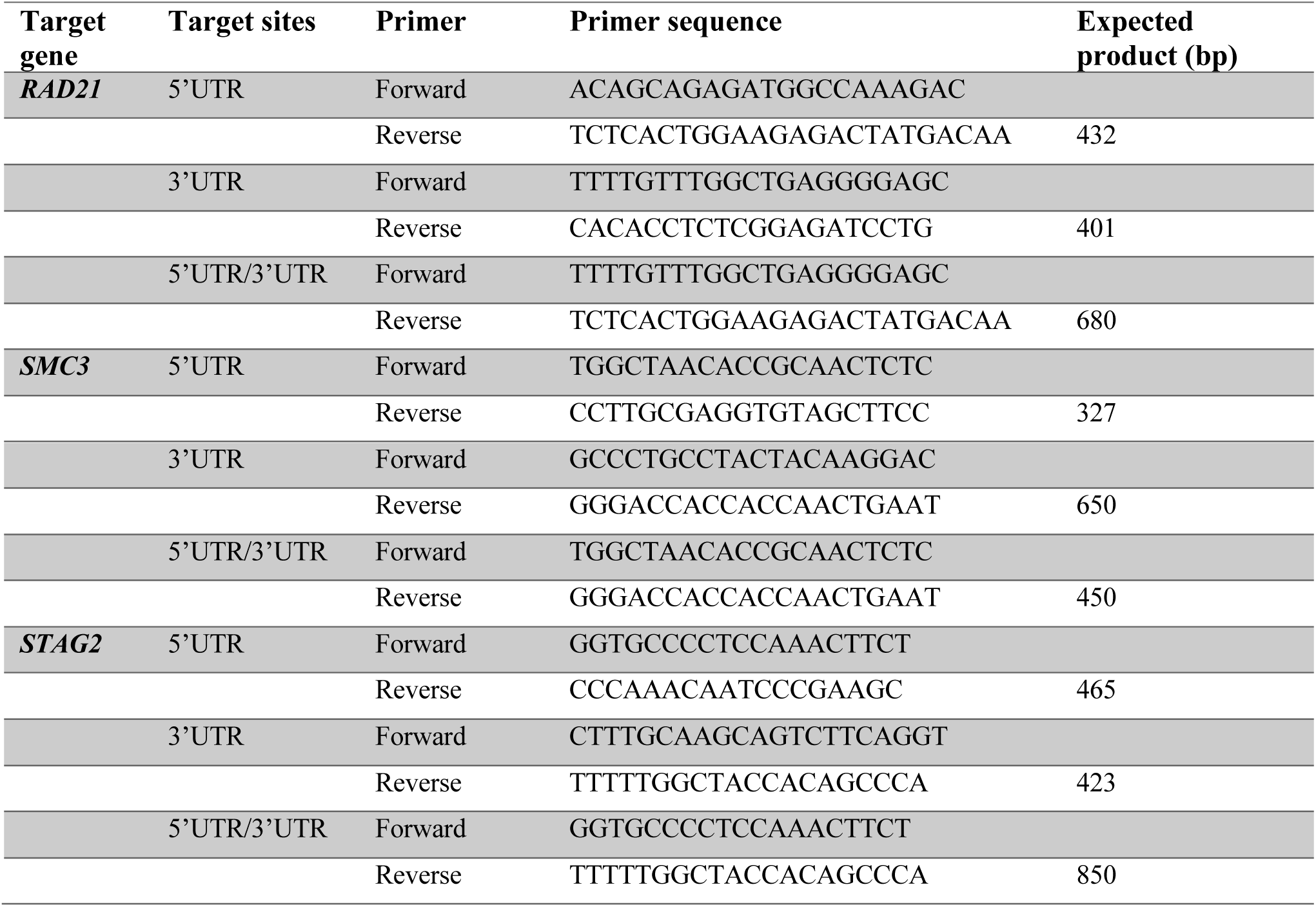
Primer sequences used in PCR assay to screen for cells with heterozygous deletion of RAD21 or SMC3 and homozygous deletion of STAG2.

**Supplementary Table 5.**
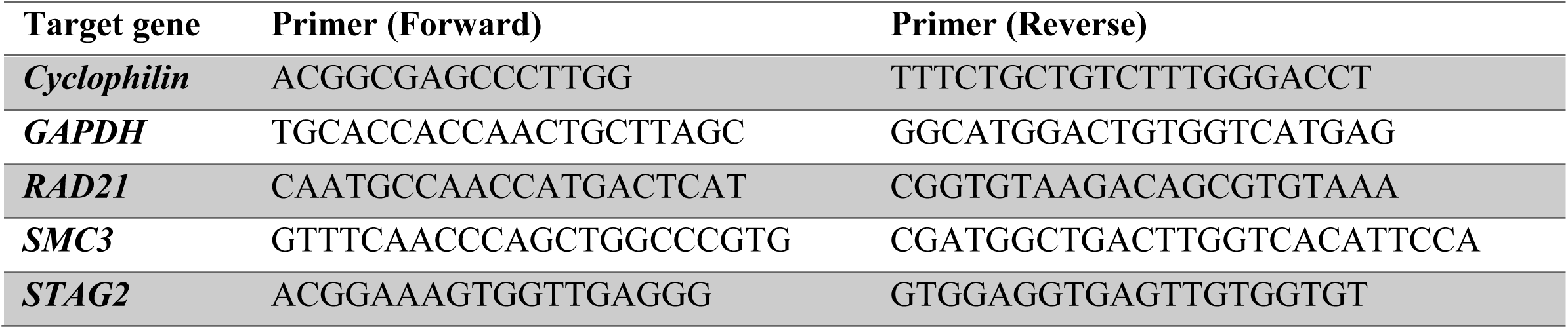
Primer sequences used in RT-qPCR.

